# The *DNAJB1–PRKACA* Oncogenic Fusion Drives Stepwise Biliary and Pancreatic Carcinogenesis from Intraductal Precursor Lesions

**DOI:** 10.64898/2026.05.23.727425

**Authors:** Patrick R. Carney, Manabu Nukaya, Jason A. Carter, Anthony J. Veltri, Kristina A. Matkowskyj, Austin Stram, C. Dustin Rubinstein, Alex C. Veith, Venu G. Pillarisetty, Christopher A. Bradfield, Sean M. Ronnekleiv-Kelly

**Author notes:** **Corresponding Author:** Sean M. Ronnekleiv-Kelly, MD, Address: 600 Highland Ave. Bx 7375, K3/705 CSC, Madison, WI, 53792.

## Abstract

Intraductal oncocytic papillary neoplasms (IOPNs) are rare tumors that develop from biliary and pancreatic ductal epithelium and progress to lethal cancers. Human IOPNs and IOPN-associated carcinomas are known to harbor the *DNAJB1-PRKACA* gene fusion, however the role of the oncogenic fusion in carcinogenesis is poorly understood. We developed a Cre-inducible mouse model of human DNAJB1-PRKACA expression and substantiated that the *DNAJB1-PRKACA* fusion gene is a bona fide driver of IOPN-associated biliary and pancreatic carcinoma. By analyzing the unique histopathologic and transcriptional changes that occur at each stage of tumor development, we found that these murine tumors closely mimic human *DNAJB1-PRKACA* driven tumors. Furthermore, we identified that many of the salient features of invasive carcinoma are established at the pre-invasive stage, including evidence of tumor cell metabolic dysregulation, immunosuppressive tumor stroma and expression of genes strongly associated with invasive *DNAJB1-PRKACA* driven cancer in humans. Finally, we found that *Slc16a14*, a characteristic *DNAJB1-PRKACA* regulated super-enhancer associated gene, serves as a robust biomarker of malignant transformation from IOPN to invasive carcinoma.

## INTRODUCTION

Intraductal oncocytic papillary neoplasms (IOPNs) are rare neoplasms arising from the biliary and pancreatic ductal epithelium that carry malignant potential, progressing to IOPN-associated cholangiocarcinoma (CCA) and pancreatic carcinoma^1, 2^. Our current understanding of these cancers is based on human case report series that suggest carcinogenesis occurs in a stepwise progression from IOPN to invasive carcinoma.^2–4^ These series also underscore that human IOPNs and IOPN-associated carcinomas have historically been distinguished from other pancreatic and bile duct tumors by their histologic appearance. Key histologic elements include arborizing papillae containing cells with abundant granular and eosinophilic (oncocytic) cytoplasm – cytologic features that reflect densely packed mitochondria suggestive of profound metabolic dysfunction.^3–5^ These tumors are also characterized by a desmoplastic stroma rich in collagen fibers surrounding carcinoma cells.^2, 6, 7^ In addition, IOPN-associated carcinomas do not harbor conventional oncogenic mutations (e.g., *KRAS*, *GNAS*, *TP53*, *CTNNB1*, *SMAD4*) classically found in other pancreaticobiliary malignancies^2, 6, 8^. Instead, IOPNs and IOPN-associated carcinomas of the bile duct and pancreas harbor the *DNAJB1-PRKACA* fusion oncogene, strongly implicating this *PRKACA* fusion as a molecular driver. ^2, 3, 5, 6, 8–11^

The *DNAJB1-PRKACA* oncogene fusion is the result of a 400 kbp deletion on chromosome 19 in humans, between DnaJ homolog subfamily B member 1 gene (*DNAJB1*) and protein kinase catalytic subunit alpha (*PRKACA*) gene.^12^ The structural rearrangement places exon 1 of *DNAJB1* in sequence with exons 2-10 of *PRKACA*. Loss of exon 1 in *PRKACA* results in a chimeric protein (DNAJ-PKAc) that is no longer bound by PKA regulatory subunits, and consequently, the normally cAMP responsive protein demonstrates unbridled PKA activity, causing cellular transformation. Until recently, *DNAJB1-PRKACA* was thought to be highly specific for another rare liver cancer, fibrolamellar carcinoma (FLC)^12–15^. FLC is not only driven by the same fusion oncogene, but also displays strongly overlapping histologic features including large oncocytic cells with abundant granular cytoplasm containing densely packed mitochondria, akin to the observations in IOPN-associated carcinomas.^16–18^ Further evidence of biologic convergence of these two disease processes comes from the fact that aberrant Protein Kinase A (PKA) activity is central to tumorigenesis.^19–22^ Transcriptional analysis of *DNAJB1-PRKACA* expressing tumors of the liver and pancreatic ductal cells cluster transcriptionally with FLC.^23^ These observations suggest these distinct rare and lethal cancers may in fact be biologically synonymous.

Admittedly, authors in each of the series investigating IOPN-associated carcinoma acknowledged that the tumorigenesis of IOPN-associated carcinoma remains unclear.^24–27^ The authors further conceded that many unanswered questions exist and are exceedingly challenging to answer absent a model system (due to the rare nature of this lethal cancer). For instance, advanced pre-cancerous tumors or invasive carcinoma have been identified in humans^2, 4, 6^, but the carcinogenic process remains unclear. Understanding how these carcinomas develop and progress would enable key insights that should lead to biomarkers of detection and novel therapeutic targets in *DNAJB1-PRKACA*-driven cancer. Concordantly, this is the same challenge faced in FLC (also a rare and lethal cancer driven by *DNAJB1-PRKACA*), where there are currently no model systems available to investigate the pathogenesis of tumor initiation and progression in the native microenvironment.

We capitalized on this concept to overcome the barrier of lack of knowledge of IOPN-associated carcinoma development. We placed the *DNAJB1-PRKACA* gene fusion into a safe harbor locus in mice to allow control of cell specificity and timing of oncogene fusion activation. Cell-specific expression enabled us to direct *DNAJB1-PRKACA* activation to bile duct and pancreatic duct cells, which are the cells of origin for IOPN-associated cholangiocarcinoma and IOPN-associated pancreatic carcinoma, respectively. Consequently, we present here the carcinogenic process of these tumor types.

## RESULTS

### Constructing a conditional in vivo model of DNAJB1-PRKACA expression

Expression of the *DNAJB1-PRKACA* fusion oncogene has been identified in three distinct human cancers; FLC, IOPN-associated CCA and IOPN-associated pancreatic carcinoma.^2, 6, 8, 9, 12^ Compelling data have revealed that the *DNAJB1-PRKACA* fusion is the primary driver of FLC,^12–14^ but the same level of evidence is not present for the IOPN-associated carcinomas. To test the hypothesis that *DNAJB1-PRKACA* is similarly a driver of IOPN-associated carcinomas, we constructed a genetically engineered mouse model system harboring conditional expression of the fusion oncogene. This would allow targeting of *DNAJB1-PRKACA* expression to biliary and pancreatic ductal epithelium, which together are thought to represent the cells of origin for IOPNs and associated carcinomas. Further, our strategy employed fluorescent marker tagging of *DNAJB1-PRKACA* expressing cells to facilitate reliable tracking of tumor development.

To accomplish this, we used CRISPR/Cas9 to insert human *DNAJB1–PRKACA* cDNA followed by an internal ribosomal entry site (*I*, *IRES*) in the *Ai14* red fluorescent protein reporter mouse (*Rosa26^CAG-LSL-tdTomato^*). This placed the *DNAJB1-PRKACA* fusion cDNA and *IRES* in sequence with tdTomato red fluorescent protein at the *Rosa26* safe harbor locus to generate the *Ai14^D-P^* mouse (*Rosa26^CAG-LSL-DNAJB^*^1^*^-PRKACA-IRES-tdTomato^*). Long-read whole genome sequencing, restriction enzyme mapping and PCR genotyping confirmed integration of the *DNAJB1-PRKACA*-IRES cassette between the *LoxP-Stop-LoxP* and tdTomato construct sites in Ai14 mice (**Figure 1A-C**). To confirm that (1) human DNAJB1-PRKACA protein (DNAJ-PKAc) and (2) tdTomato red fluorescent protein expression occurred under the control of Cre recombinase, we injected *Ai14^D-P^* mice with a liver tropic adeno-associated virus variant 8 (AAV8) containing CMV promoter driving Cre recombinase expression vector (AAV8-CMV-Cre) or no Cre expression vector (AAV8-Null). Two weeks after AAV8 injection, livers were isolated from the injected *Ai14^D-P^* mice, and DNAJ-PKAc and tdTomato protein expression were analyzed. This confirmed DNAJ-PKAc and tdTomato protein expression in the AAV8-CMV-Cre injected mice **(Figure 1D)**. Of note, DNAJ-PKAc protein is a larger protein than native PKAc,^12^ and only native PKAc was detected in the AAV8-Null injected mice. Furthermore, tdTomato protein expression was confirmed in liver sections of mice treated with AAV8-CMV-Cre vector using confocal fluorescence microscopy **(Figure 1E)**.

**Figure 1.**
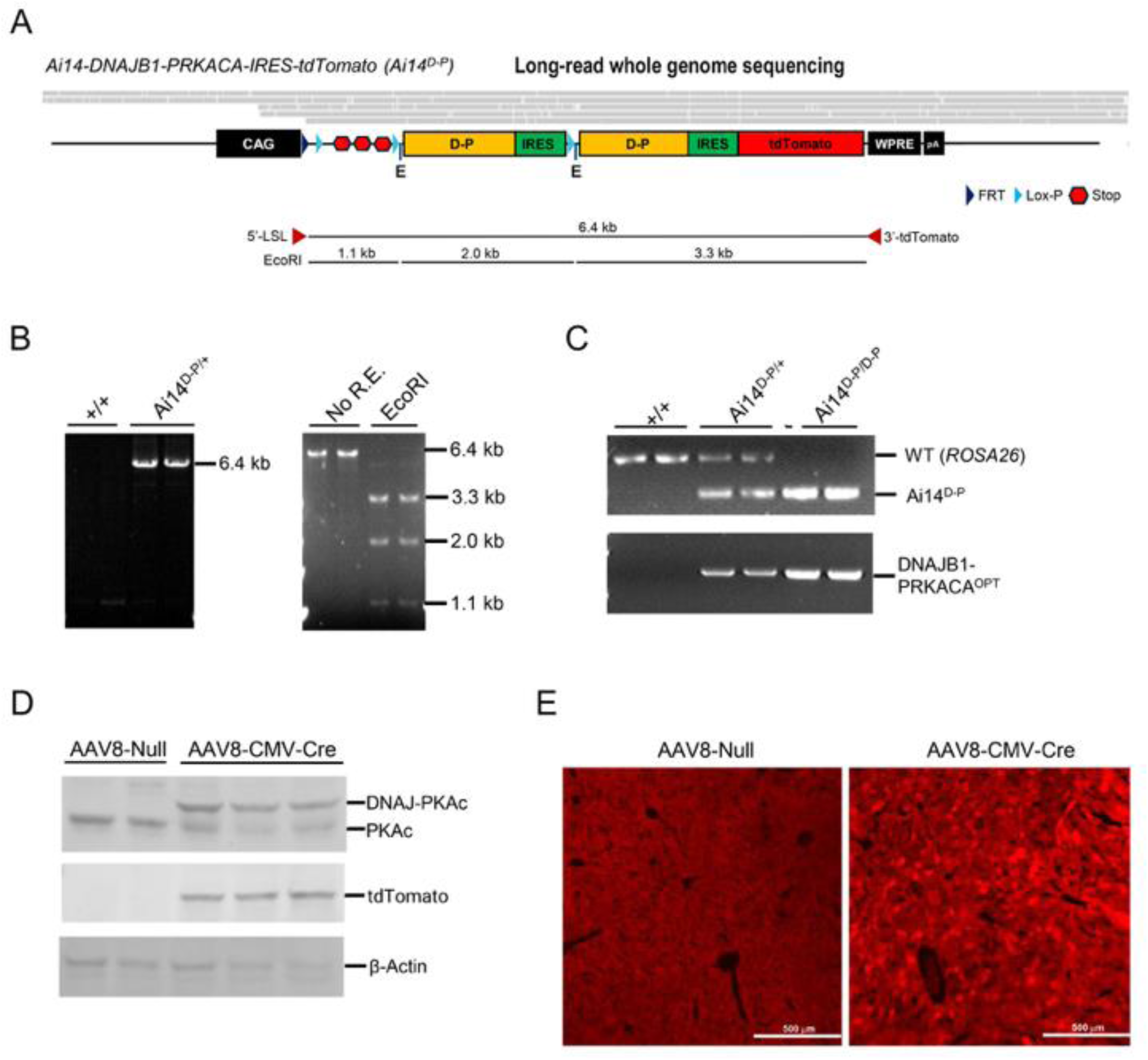
Generation of human *DNAJB1-PRKACA* expression- fluorescent reporter mouse model. (**A**) Genomic structure of Ai14^D-P^ (*Rosa26^CAG-LSL-hDNAJB1-PRKACA-IRES-tdTomato^*) mouse model. The Ai14^D-P^ allele was created by modifying the Ai14 allele *(B6.Cg-Gt(ROSA)26Sor^tm14(CAG-tdTomato)Hze^/J)*. The *hDNAJB1-PRKACA*-*IRES* and *hDNAJB1-PRKACA*-*IRES*-*tdTomato* sequences were inserted downstream of the *loxp-stop-loxp* cassette. The genomic sequence was confirmed using Oxford Nanopore long-read whole-genome sequencing. Grey bars represent full length reads spanning the Ai14^D-P^ allele, aligned to the depicted sequence. E: EcoRI site. (**B**) Gene screening using PCR. Left: The 6.4 kb sequence, including *loxp-stop-loxp*, *D-P-IRES*, and *D-P-IRES-tdTomato* cassettes, was analyzed using PCR in genomic DNA isolated from tail of WT (^+/+^) and Ai14^D-P^ (Ai14^D-P/+^) mice. The 6.4 kb amplified band was detected in Ai14^D-P^ mice. Right: The 6.4 kb amplified band was digested by EcoRI restriction enzyme. The 6.4 kb band was digested into 1.1, 2.0, and 3.3 kb fragments, respectively. (**C**) PCR genotyping for WT and Ai14^D-P^ alleles. Upper: The 297 bp (ROSA26S) and 196 bp (*tdTomato-WPRE*) amplified bands were generated from WT (^+/+^) and Ai14^D-P^ (Ai14^D-P/+^, Ai14^D-P/D-P^) alleles, respectively. Lower: The 362 bp (codon-optimized *hDNAJB1-PRKACA*) bands were amplified from Ai14^D-P^ alleles (Ai14^D-P/+^, Ai14^D-P/D-P^). (**D**) Western blot depicts the analysis for DNAJ-PKAc (*hDNAJB1-PRKACA* encodes for DNAJ-PKAc) and tdTomato protein expression. DNAJ-PKAc and tdTomato proteins were detected using liver homogenate (40 μg) from AAV8-null or AAV8-CMV-Cre injected Ai14 D-P (Ai14^D-P/D-P^) mice. DNAJ-PKAc and native PKAc protein was detected using anti-PKAc antibody. β-actin was used as a housekeeping control. The samples were loaded separately and membranes were developed independently. (**E**) Analysis of tdTomato-positive cells. The tdTomato-positive cells (red) were detected in liver from AAV8-null or AAV8-CMV-Cre injected Ai14 D-P (Ai14^D-P/D-P^) mice using a confocal microscope.

Additional validation was performed by isolating tdTomato-positive cells (tdTomato+) and tdTomato-negative cells (tdTomato-) from dissociated liver of *Ai14^D-P^*; *Krt19^CreERT^* mice **(Supplemental Figure 1D).** From the long-read WGS and restriction enzyme mapping, we found that the *Ai14^D-P^* mice carry a duplicate *DNAJB1-PRKACA-IRES* cassette between the *LoxP-Stop-LoxP* and *DNAJB1-PRKACA*-*IRES*-*tdTomato* constructs (**Figure 1A-B**). These concatemeric insertions occur frequently in targeted genetic knockins.^28, 29^ In the tdTomato-expressing cells (tdTomato+) isolated from the liver of *Ai14^D-P^; K19^CreERT^* mice, two types of Cre-*LoxP* mediated excision were observed (**Supplemental Figure 1A-B**). One excised only the Stop cassette (excision 1) and another excised both the Stop and the extra *DNAJB1-PRKACA-IRES* cassettes floxed by *Lox-P* sites (excision 2). Liver cells harboring either excised construct expressed both DNAJ-PKAc and tdTomato proteins (**Supplemental Figure 1C-D**), yielding reliable expression of the fusion oncoprotein and tdTomato protein in *Ai14^D-P^*mice.

### *DNAJB1-PRKACA* expression causes neoplastic change in biliary and pancreatic ductal epithelium

Cytokeratin 19 is a protein encoded by the *Krt19* gene, which is robustly expressed in both bile duct epithelium and pancreatic duct epithelium.^30^ Further, *Krt19* is not expressed in hepatocytes or pancreatic acinar cells. Accordingly, we selected the *Krt19^CreERT^* mouse, in which tamoxifen-inducible Cre-recombinase (Cre:ERT) is expressed in *Krt19*-expressing cells. This allows for control of both cell specificity and timing of Cre-recombinase expression in these target tissues.^30^ Regulation of timing is achieved by tamoxifen administration, which allows for translocation of Cre-recombinase to the nucleus and recombination (i.e., excision) between *Lox-P* sites. Control of fusion oncogene expression timing was an important consideration because published data support that onset of *DNAJB1-PRKACA* is a somatic event and not a germline alteration.^12^

*Krt19^CreERT^* mice were crossed to *Ai14^D-P^* mice to generate the *Ai14^D-P^; K19^CreERT^* mouse (**Figure 2A**). We administered tamoxifen to *Ai14^D-P^; K19^CreERT^* mice at age 6-8 weeks via intraperitoneal (IP) injection to test whether *DNAJB1–PRKACA* fusion gene activation (and concomitant tdTomato expression) occurred in biliary and pancreatic epithelium. In these mice, administration of tamoxifen resulted in tumor initiation characterized by the onset of *DNAJB1-PRKACA* expression, followed by progressive neoplastic changes, IOPN development and ultimately carcinoma formation by 5 months (timeline in **Figure 2B-C**). This occurred in both liver and pancreas. Focusing initially on the early neoplastic changes caused by *DNAJB1-PRKACA*, we found that within one month of tamoxifen administration there was evidence of cellular transformation / tumor initiation of the bile duct epithelium and the pancreatic duct epithelium. Tumor initiation led to development of early eosinophilic neoplasms that occurred within 1-2 months after tamoxifen administration (**Figure 2B-C**). Due to the genetic construct of these mice (*DNAJB1-PRKACA-IRES-tdTomato*), neoplastic *DNAJB1-PRKACA* expressing cells concomitantly express tdTomato protein. The precise overlap of eosinophilic cells and tdTomato+ cells was apparent on consecutive sections of Hematoxylin & Eosin (H&E) and immunohistochemical (IHC) stained slide with antibody against tdTomato protein. Notably, the cell morphology that we identified in these earliest lesions closely mimics the cell morphology of human *DNAJB1-PRKACA*-driven carcinomas, including pathognomonic cytologic features such as enlarged nuclei and abundant granular and eosinophilic cytoplasm indicative of densely packed mitochondria.^5, 6, 16–18, 31–33^ Therefore, we established that *DNAJB1-PRKACA* expression (DNAJ-PKAc) and tdTomato expression occurred in the target tissue (biliary duct and pancreatic duct epithelium), and that the early cell morphology mirrors human carcinoma cell morphology.

**Figure 2.**
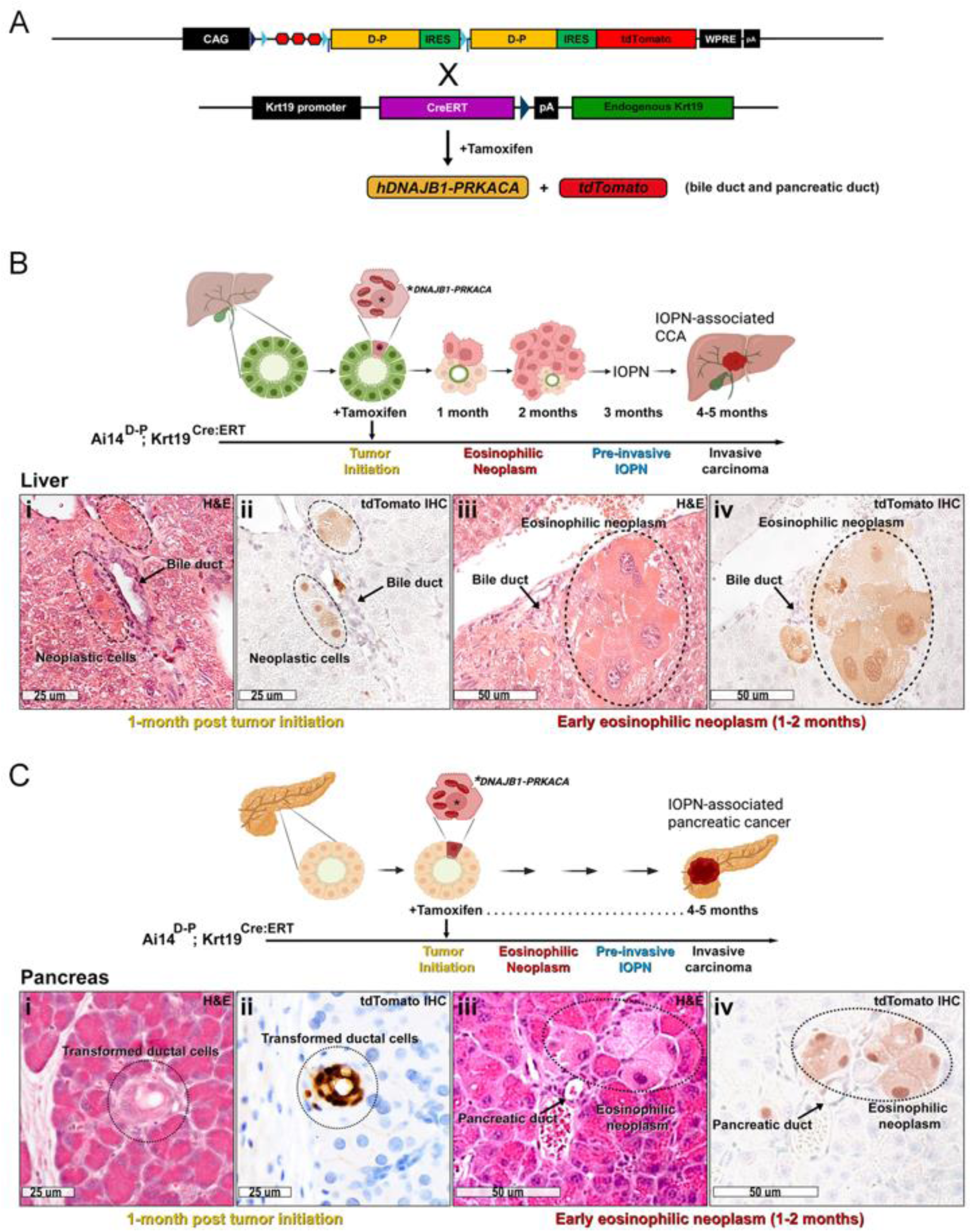
Tumor initiation by *DNAJB1-PRKACA* in biliary duct and pancreatic duct cells. (**A**) Crossing Ai14^D-P^ mice (*Rosa26^CAG-LSL-DNAJB1-PRKACA-IRES-tdTomato^*) to *Krt19^Cre:ERT^* mice results in Cre:ERT expression in *Krt19* expressing cells, including bile duct and pancreatic duct. Administration of tamoxifen results in Cre translocation to the nucleus and Cre mediated excision of *loxP-stop-loxP* sequence in these cells, resulting in expression of *hDNAJB1-PRKACA* and *tdTomato*. (**B**) Tamoxifen administration results in tumor initiation (expression of *DNAJB1-PRKACA*) in bile duct cells within the liver. This transformed cell evolves into neoplastic eosinophilic cells within 1-2 months of tamoxifen, and subsequently into a pre-invasive IOPN by 3 months. This ultimately progresses to invasive cholangiocarcinoma by 4-5 months. (**i-ii**) Shown is Hemotoxylin & Eosin (H&E) and IHC (anti-RFP) of mouse liver from *Ai14^D-P^; Krt19^Cre:ERT^* mice at 1-month after tamoxifen administration. This captures the earliest development of neoplastic cells (black ellipses) adjacent to a bile duct (arrow). The eosinophilic cells show tdTomato positivity via IHC. (**iii-iv**) Following initiation, multicellular eosinophilic neoplasms develop within 1-2 months consisting of characteristic eosinophilic cells with abundant granular cytoplasm and enlarged nuclei (black ellipse) adjacent to a bile duct (arrow). The eosinophilic cells transformed by *DNAJB1-PRKACA* express tdTomato as seen on IHC. (**C**) Shown is a parallel progression in the pancreas with pancreatic duct cell tumor initiation occurring at the time of tamoxifen administration (*DNAJB1-PRKACA* induced transformation). Eosinophilic cells progress to eosinophilic neoplasms, IOPN and IOPN-associated pancreas carcinoma. (**i-ii**) Shown is H&E and IHC demonstrating the earliest onset of cellular transformation (black circle) involving a pancreatic duct. (**iii-iv**) By 1-2 months after tamoxifen administration, early eosinophilic neoplasms form (black ellipse) adjacent to a pancreatic duct (black arrow) where the eosinophilic cells concomitantly express tdTomato as seen on IHC.

### *DNAJB1-PRKACA* driven carcinomas are lethal

In contrast to the early microscopic lesions visualized shortly after tumor initiation, there are relatively large cystic and solid tumors evident in the pancreas and liver by ∼4-5 months post tamoxifen induction. For example, pre-invasive pancreatic IOPNs strongly expressing tdTomato protein (and likewise DNAJ-PKAc) appear as discrete lesions that are well visualized (black arrow) in the background of a relatively normal-appearing pancreas (**Figure 3A**). As the IOPNs become more complex, the surrounding normal pancreatic tissue is further displaced as the cystic tumors predominate (**Figure 3B**). These pre-invasive pancreatic IOPNs progress to invasive carcinoma, where the pancreas becomes markedly distorted and fibrotic adjacent to the invasive IOPN-associated pancreatic carcinomas (**Figure 3C**). In parallel, pre-invasive biliary IOPNs and IOPN-associated cholangiocarcinoma (CCA) form in the liver. The pre- invasive IOPNs that arise from bile duct are cystic in nature (**Figure 3D-E**), and the transition to invasive IOPN-associated carcinoma is evident by a more solid appearing mass (**Figure 3F**). As the invasive carcinomas progress, they ultimately become life-limiting. For example, the carcinomas invade surrounding structures such as abdominal wall (**Figure 3G**) or large vessels (**Figure 3I**), and the primary tumor cells as well as the vessel-invasive tumor cells are strongly positive for tdTomato expression (**Figure 3H**, **3J**). Survival analysis by Kaplan-Meier (**Figure 3K**) demonstrates that median survival for these mice is approximately 115 days after tamoxifen induction as compared to normal lifespan of control mice (log-rank *p* < 0.0001). Interestingly, outside of the IOPN-associated CCA and pancreatic carcinoma, separate soft tissue carcinomas of the limbs also develop in parallel with the biliary and pancreatic tumors (**Supplemental Figure 2**). These soft tissue tumors arise from *Krt19*-expressing cells in the sweat glands (tdTomato+, cytokeratin+) and are reminiscent of human malignant sweat gland tumors on review by a board-certified pathologist. The malignant nature of these tumors was confirmed on histology. Further validation of malignancy was demonstrated by dissociating the soft tissue tumors and reimplanting the tumor cells into the flank of C57Bl/6J wild-type mice, which grew into identical appearing soft tissue tumors (**Supplemental Figure 2**). Notably, the histologic features of these murine soft tissue tumors are consistent with *DNAJB1-PRKACA*-driven cancer, including dense fibrous bands and enlarged eosinophilic tumor cells. Collectively, the *DNAJB1-PRKACA* tumor model system progresses over a relatively short duration despite arising in an immune competent mouse, strongly indicating that *DNAJB1-PRKACA* promotes an immune suppressed microenvironment. This coincides with the immunosuppressive tumor microenvironment seen in human *DNAJB1-PRKACA* driven carcinoma (e.g., FLC).^34–36^ Furthermore, development of invasive carcinoma is not protracted which is valuable for evaluation of carcinogenesis and treatment response.

**Figure 3.**
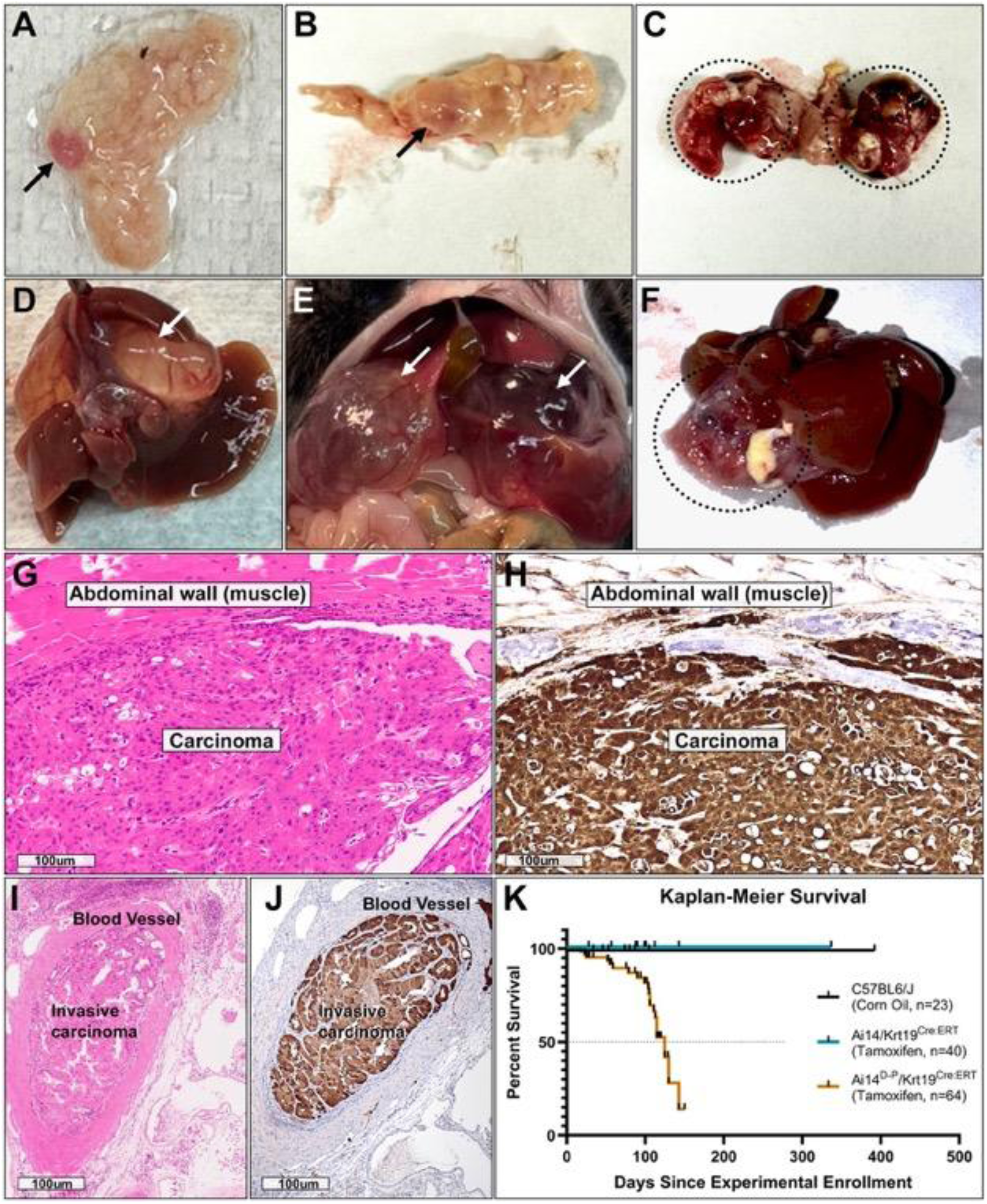
*DNAJB1-PRKACA* induced IOPNs progress to lethal invasive IOPN-associated carcinomas. Ai14^D-P^ mice crossed to *Krt19-Cre:ERT* mice are administered tamoxifen to activate the *DNAJB1-PRKACA* allele in ductal cells. **(A-C)** Images of tumor in pancreas. Shown is an IOPN lesion (arrow) arising in the background of relatively normal and translucent pancreas (**A**). This progresses to a larger pre-invasive IOPN (arrow) where the associated pancreas becomes more fibrotic (**B**). Eventually, invasive carcinoma forms, with associated markedly abnormal pancreas including significant fibrosis / desmoplasia (black circles) (**C**). (**D-F**) A parallel progression occurs in the liver where pre-invasive neoplasms form (white arrows) that progresses to an invasive carcinoma, causing significant deformity and fibrosis in the liver (black circle). (**G-J**) The invasive carcinomas ultimately grow into adjacent structures, such as abdominal wall, and demonstrate metastatic spread including lymphovascular invasion. Shown is H&E (**G, I**) and tdTomato IHC (**H, J**) of invasive tumor growing into abdominal wall and into a blood vessel. (**K**) Shown is a Kaplan-Meier survival curve of control mice (C57BL6/J and *Ai14; Krt19^Cre:ERT^*) versus study mice (*Ai14^D-P^*; *Krt19^Cre:ERT^*). No control mice died during the time evaluated, and liver / pancreas histology was normal. Meanwhile, study mice harboring *DNAJB1-PRKACA* manifested liver and pancreas pathology that was ultimately life limiting. On average, study mice survived ∼4-5 months following activation of *DNAJB1-PRKACA*.

### *DNAJB1-PRKACA* drives IOPN-associated cholangiocarcinoma

We identified nascent eosinophilic neoplasms in the *Ai14^D-P^; Krt19^CreERT^* mouse liver at 1-2 months (**Figure 2B**) and found that these *DNAJB1-PRKACA*-driven tumors progress to macroscopically evident invasive carcinomas by roughly five months after tamoxifen induction (**Figure 3D-F**). Between one month and five months, there is a remarkable and systematic progression of the neoplastic cells caused by *DNAJB1-PRKACA*. By approximately two months following tamoxifen administration, larger eosinophilic neoplastic cell clusters emerge (**Figure 4A-D**). Like the earliest lesions, the neoplasms arise adjacent to bile ducts and the neoplastic cells demonstrate abundant eosinophilic cytoplasm with enlarged nuclei (**Figure 4A**), which is readily apparent on magnified views (**Figure 4C**). The tumor cells are strongly positive for tdTomato (**Figure 4B**, **4D**), substantiating the genetic construct that promotes fusion oncoprotein expression concomitant with tdTomato protein expression. By three to four months, the eosinophilic neoplasms progress to oncocytic papillary neoplasms (IOPNs) that closely resemble human lesions.^8^ The early pre-invasive IOPNs harbor increasingly complex architecture (**Figure 4E-F**), and are comprised of characteristic neoplastic eosinophilic cells lining the cystic structures with papillary projections (**Figure 4G-H**). The early IOPNs grow into advanced IOPNs which are distinguished by development of a dense intratumoral stroma (**Figure 4I-J**). On closer examination, these complex lesions have clear evidence of significant fibrosis between the eosinophilic (and tdTomato+) neoplastic cells (**Figure 4K-L**). The fibrotic stroma is a defining feature of *DNAJB1-PRKACA* driven cancers,^16^ and this fibrosis is thought to contribute to the marked immunosuppressive tumor microenvironment of *DNAJB1-PRKACA*-driven carcinoma.^37^ Consistent with this idea, the advanced IOPNs progress to invasive IOPN-associated CCA, where cancer cells invade the surrounding stroma adjacent to the neoplastic IOPN cells that line dilated cystic spaces (**Figure 4M**). Magnified views demonstrate that the carcinoma contains discohesive cells with disorganized growth and irregular nuclei (**Figure 4Q**). Reflecting the cell of origin, the neoplastic cells and carcinoma cells express Cytokeratin 19 protein (**Figure 4N**) and the Cytokeratin positive carcinoma cells are present amidst the desmoplastic stroma (**Figure 4R**). The desmoplastic stroma of the cancer microenvironment is highlighted by wispy blue-grey stroma on trichrome stain (**Figure 4O**, **4S**), and the neoplastic cells and carcinoma cells are easily depicted by tdTomato protein expression (**Figure 4P**, **4T**). Collectively, we confirm that onset of *DNAJB1-PRKACA* in biliary epithelium results in characteristic eosinophilic cells that progress to pre-cancerous IOPNs and ultimately to invasive carcinoma over a span of five months after tamoxifen induction. Notably, the advanced pre-malignant IOPNs and invasive carcinomas displayed a dense fibrous stroma, which is a feature that has not been previously established in immune competent models of *DNAJB1-PRKACA* associated neoplasms. Also of critical impact, we have established a *DNAJB1-PRKACA* driven IOPN-associated CCA *in vivo* model, and this model substantiates the human case-series IOPN-associated carcinoma data.

**Figure 4.**
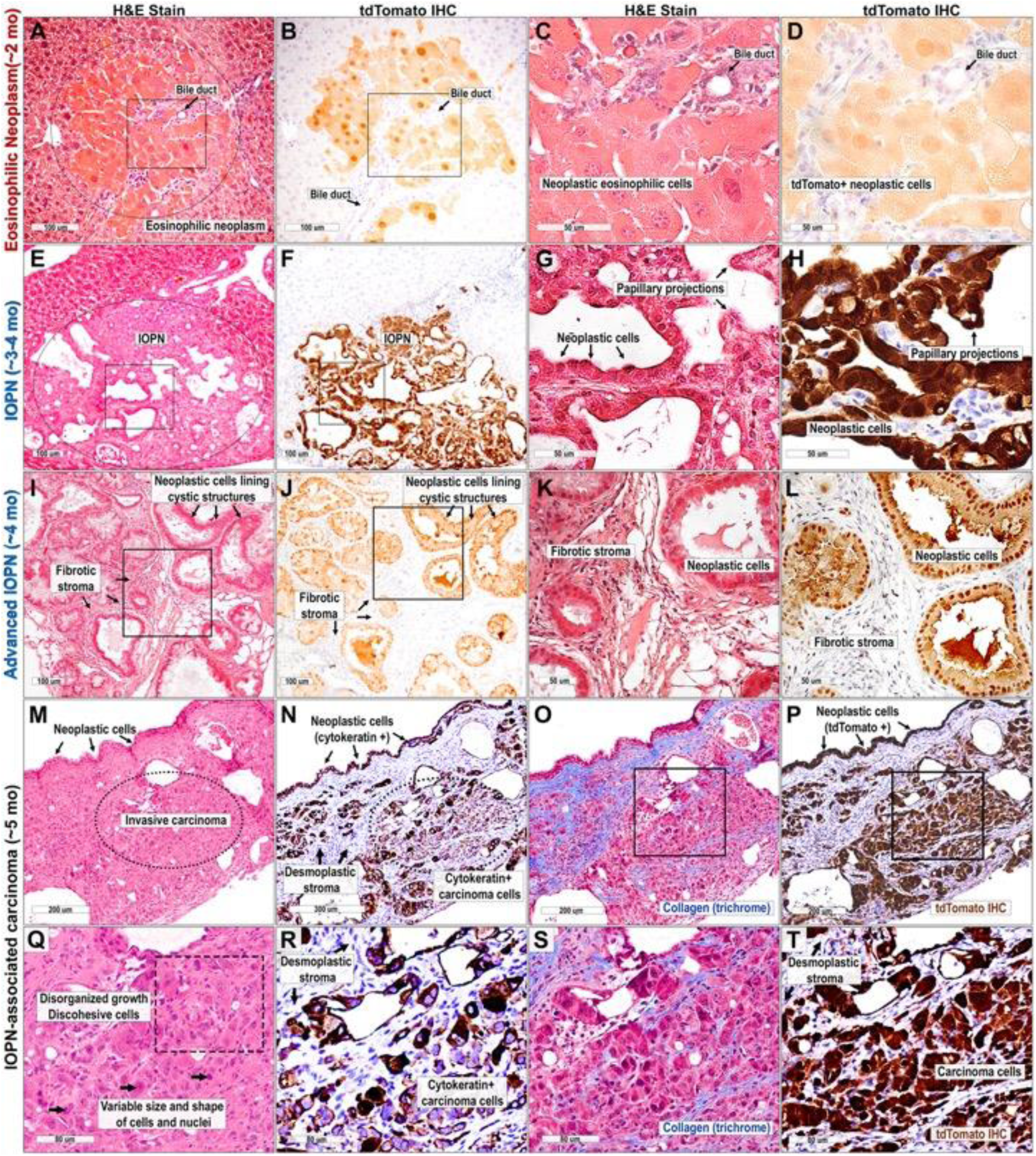
*DNAJB1-PRKACA* drives a progression from eosinophilic neoplasm to IOPN-associated cholangiocarcinoma. (**A-D**) Shown is H&E and IHC (tdTomato) of an eosinophilic neoplasm. **(A-B)** Activation of the *DNAJB1-PRKACA* oncogene in *Krt19* expressing cells (bile duct) results in formation of characteristic neoplasms containing large eosinophilic cells with abundant pink granular cytoplasm and large nuclei (black circle) adjacent to / arising from the bile duct. (**C-D**) Magnification views of the regions the black box from A and B. These oncocytic cells appear histologically consistent with transformed cells and reflect densely packed mitochondria, similar to cytologic features seen in Fibrolamellar cancer. The transformed oncocytic cells are strongly positive for tdTomato, which serves as a marker for *DNAJB1-PRKACA* expression (i.e., *DNAJB1-PRKACA-Ires-tdTomato*). **(E-F)** The eosinophilic neoplasms (Eos) progress to cystic neoplastic lesions consistent with intraductal oncocytic papillary neoplasms (IOPN) which demonstrate cystic structures containing papillary projections that are tdTomato positive. **(G-H)** Magnified views (from region in black box) demonstrate that the IOPNs manifest papillae with *DNAJB1-PRKACA* expressing neoplastic cells lining the cystic structures (tdTomato+). **(I-J)**. The pre-cancerous cystic neoplasms progress to advanced cystic neoplasms harboring oncocytic cells and demonstrate clear evidence of significant fibrosis surrounding the neoplastic cells. This fibrosis is more clearly seen on magnification views (**K-L**, representing the region in the black box) and is an essential and defining feature of tumors caused by *DNAJB1-PRKACA*. **(M-P)** By age 4-5 months following activation of *DNAJB1-PRKACA*, these mice develop invasive cholangiocarcinoma. Carcinoma cells are positive for cytokeratin indicating epithelial origin and trichrome stain reveals collagenous bands as well as desmoplastic stroma within the tumor. **(Q-T)** On higher magnification, the carcinoma cells are observed to be discohesive and disorganized (**Q**). The cancer cells demonstrate positive cytokeratin (**R**), collagenous bands with desmoplasia (**S**) and demonstrate strong tdTomato positivity (**T**), indicating robust *DNAJB1-PRKACA* expression.

### *DNAJB1-PRKACA* drives IOPN-associated pancreatic carcinoma

Similar to the progression seen in liver, we identified that oncocytic / eosinophilic cells in the pancreas (**Figure 2C**) progress to form eosinophilic neoplasms (**Figure 5A**). Thus, this oncocytic cellular morphology is the initial histologic indication of dysregulated PKA by *DNAJB1-PRKACA* in both liver (bile duct) and pancreas. Mirroring our liver observations, these eosinophilic neoplasms progress to cystic neoplasms (early IOPNs) with papillary projections and neoplastic oncocytic cells lining the cystic structures (**Figure 5B-D**). Interestingly, we captured what appeared to be the development of an IOPN from a large eosinophilic neoplasm, demonstrating the transition from the characteristic eosinophilic neoplasms to the pre-cancerous IOPNs (**Figure 5B**). Concordant with the bile duct tumors (**Figure 4**), the pancreatic neoplastic cells consistently express tdTomato protein (**Figure 5C-D**), again validating the genetic construct of the model (*DNAJB1-PRKACA-IRES-tdTomato*). The transition between early IOPNs and more advanced IOPNs is marked by the development of a fibrous stroma (**Figure 5E-F**) and subsequently a highly complex papillary architecture of the neoplastic tumor cells (**Figure 5G**). The stroma and complex neoplastic architecture of these tumors co-develop within the tumor microenvironment, indicating robust communication between these cellular compartments (e.g., tumor cells and fibroblasts). Further, the eosinophilic proliferations and IOPN tumor cells are strongly positive for cytokeratin (**Figure 5H**), reflecting the cell of origin for these neoplasms (i.e., *Krt19* expressing cells). Reinforcing that a densely fibrotic stroma and complex neoplastic cell architecture immediately precede invasive carcinoma, we captured the precise transition from complex pre-cancerous IOPN (**Figure 5I-J**) to invasive carcinoma (**Figure 5K**) within the same neoplasm. Mimicking human *DNAJB1-PRKACA* driven cancers, we found that the tumor microenvironment is rich in collagen deposition (**Figure 5L**).^1, 2, 6, 16, 38^

**Figure 5.**
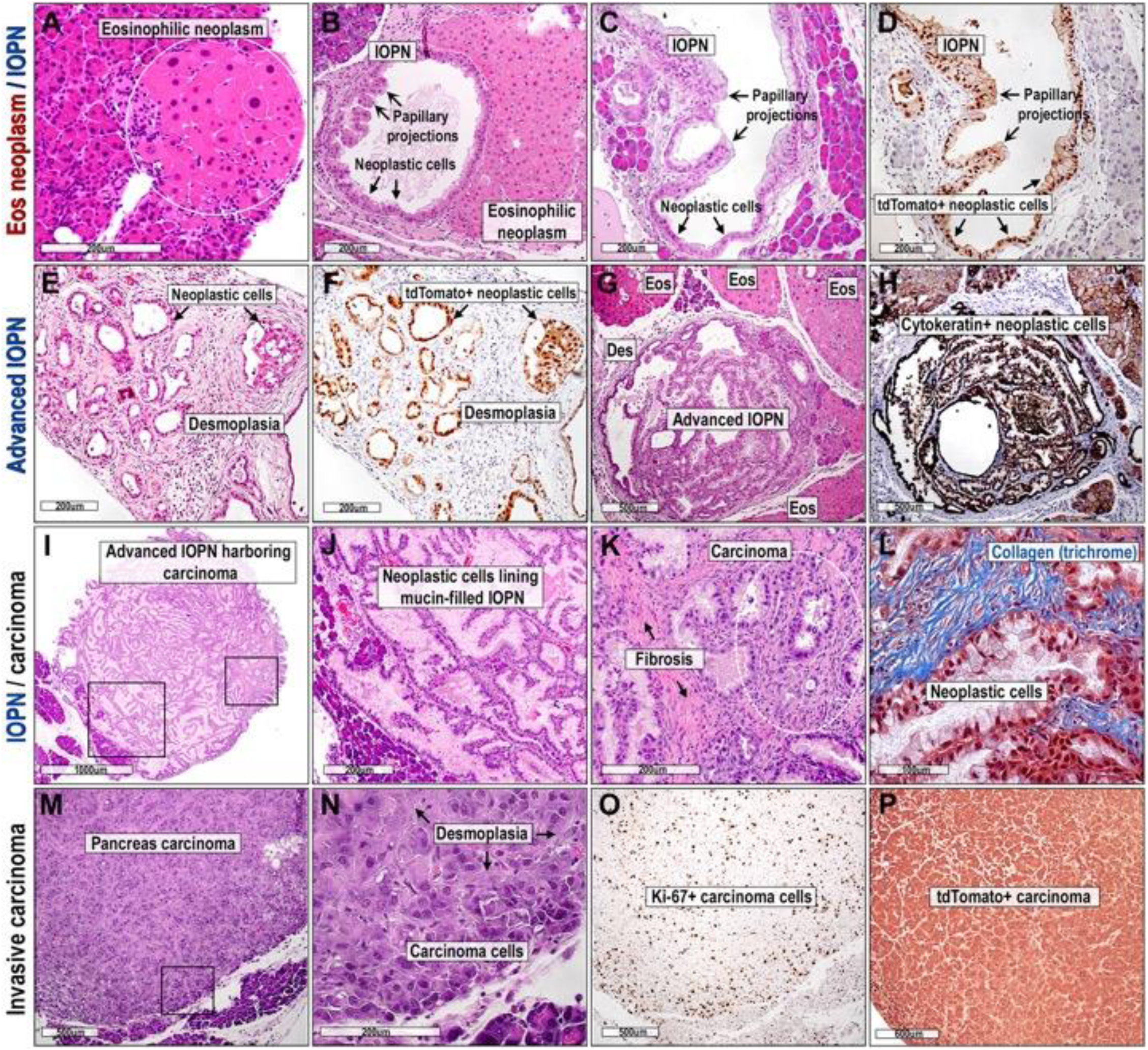
*DNAJB1-PRKACA* driven IOPN and IOPN-associated pancreatic carcinoma. **(A)** By age 4-5 weeks after tamoxifen activation of *DNAJB1-PRKACA*, mice develop cellular transformation consisting of large eosinophilic cells with abundant pink cytoplasm and large nuclei (white circle) adjacent to the pancreatic duct that appear histologically consistent with *DNAJB1-PRKACA* transformed cells. **(B)** Progression of the eosinophilic neoplastic cells yields pancreatic intraductal oncocytic cystic neoplasms with evidence of papillary projections (black arrowheads) and neoplastic cells (black arrows). (**C-D**) Shown adjacent to the H&E section is an IHC section (tdTomato) which reveals that the oncocytic cells lining the cystic neoplasms are tdTomato+ (i.e., *DNAJB1-PRKACA* expressing cells). **(E-F)** As these neoplasms progress, the architecture becomes more complex including the lining of the cystic neoplasm (arrows in **E**) and desmoplasia that co-develops around the cystic structures. IHC (tdTomato) reveals that the neoplastic cystic tumors express *DNAJB1-PRKACA* (tdTomato+), but the surrounding fibrosis does not. **(G-H)** The cystic neoplasms (IOPNs) develop significantly more complex architecture and papillary projections that are strongly positive for cytokeratin 19 (shown adjacent to H&E is cytokeratin 19 IHC), reflecting the cell of origin for these neoplasms (*Krt19^Cre:ERT^*). **(I)** Shown is an H&E of a large pancreatic IOPN tumor demonstrating the extensive papillary architecture, with boxes surrounding regions of interest. (**J**) Magnification of one region of interest (larger black box) more clearly depicts the complex architecture. (**K**) Additionally, within this IOPN, a microscopic focus of invasive carcinoma was identified (smaller black box). This region demonstrates fibrosis (pink) adjacent to the invasive carcinoma (**K**, white circle), with desmoplasia evident in the invasive carcinoma. (**L**) Intratumoral collagen deposition adjacent to neoplastic cells is highlighted by trichrome stain. **(M-N)** Ultimately, large carcinomas develop. Shown is H&E of the invasive carcinoma with extensive desmoplasia (pink). Magnification view (region in the black box) of the tumor reveals markedly atypical cellular morphology with irregular and heterogeneous nuclei. (**O**-**P**) The carcinoma cells are strongly positive for Ki-67 (**O**) and tdTomato (**P**) on IHC.

Ultimately, large invasive pancreatic carcinomas develop from the complex IOPNs (**Figure 5M**), with features that include desmoplasia interspersed between markedly irregular appearing carcinoma cells (**Figure 5N**), and evidence of increased proliferation marked by diffuse Ki-67 positivity within the tumor (**Figure 5O**). Concordant with the pre-invasive neoplastic cells caused by *DNAJB1-PRKACA* in this model, the malignant cells display robust tdTomato protein expression (**Figure 5P**). Taken together, we discovered that *DNAJB1-PRKACA* drives a highly consistent oncocytic / eosinophilic cell morphology as the first manifestation of tumor initiation, whether arising from bile duct or pancreatic duct, and this early (pre-invasive) morphology mimics the oncocytic cell morphology that is seen in (invasive) human *DNAJB1-PRKACA* driven cancer.^2, 16^ Furthermore, as DNAJ-PKAc fusion protein drives progression of the tumor cells, a characteristic fibrous stroma concomitantly develops; notably, this fibrotic stroma emerges prior to the onset of invasive carcinoma, potentially playing a role in immune evasion of tumor cells. In turn, carcinoma develops. Therefore, we show that *DNAJB1-PRKACA* oncogene fusion is sufficient as a single molecular driver of *in vivo* carcinoma formation, for both IOPN-associated CCA and IOPN-associated pancreatic carcinoma. Our mapping of initiation-promotion-progression holds significant implications for defining key contributions to *DNAJB1-PRKACA*-induced carcinogenesis.

### Co-evolution of *DNAJB1-PRKACA* fusion gene-driven neoplasms with a permissive tumor microenvironment

A unique aspect of our *DNAJB1-PRKACA* model system in comparison to prior models is the development of a highly complex tumor immune microenvironment with desmoplastic stroma.^13, 14, 18^ The fibrotic / desmoplastic tumor microenvironment is an essential feature of human *DNAJB1-PRKACA* fusion gene-driven tumors.^2, 6, 15, 16^ We hypothesized that the densely fibrotic stroma of the tumor microenvironment facilitates development of invasive carcinoma (i.e., established prior to onset of invasive carcinoma). To test this hypothesis, we performed spatial transcriptomics on a total of 15 samples for pancreatic (n = 8), liver (n = 6) and soft tissue (n = 1) tumors selected to represent the spectrum of disease ranging from early eosinophilic lesions to IOPN to invasive carcinoma (**Supplemental Figure 3**). After stringent quality control, we obtained transcriptomes from a total of 1.4 million individual cells representing these samples and comprised of 10 major cell types including parenchymal cells (e.g., hepatocytes), mesenchymal cells (e.g., cancer associated fibroblasts (CAFs)), vascular cells (e.g., endothelial), immune cells (e.g., myeloid and lymphocytes), and tumor cells (**Figure 6A-B, Supplemental Figure 4A**). These major cell types were conserved across samples (**Supplemental Figure 4B-C**) and further subdivided according to their expression of canonical marker genes (**Supplemental Figure 4D**). All major immune cell types found in human *DNAJB1-PRKACA*-driven cancer^35, 37^ were identified in these samples and there were no significant differences in the immune populations of liver and pancreatic tumors (**Supplemental Figure 5A**). Outside of organ specific cell populations (e.g., hepatocytes, acinar cells), we did not find significant differences in non-immune cell populations between liver and pancreatic tumors (**Supplemental Figure 5B**).

**Figure 6.**
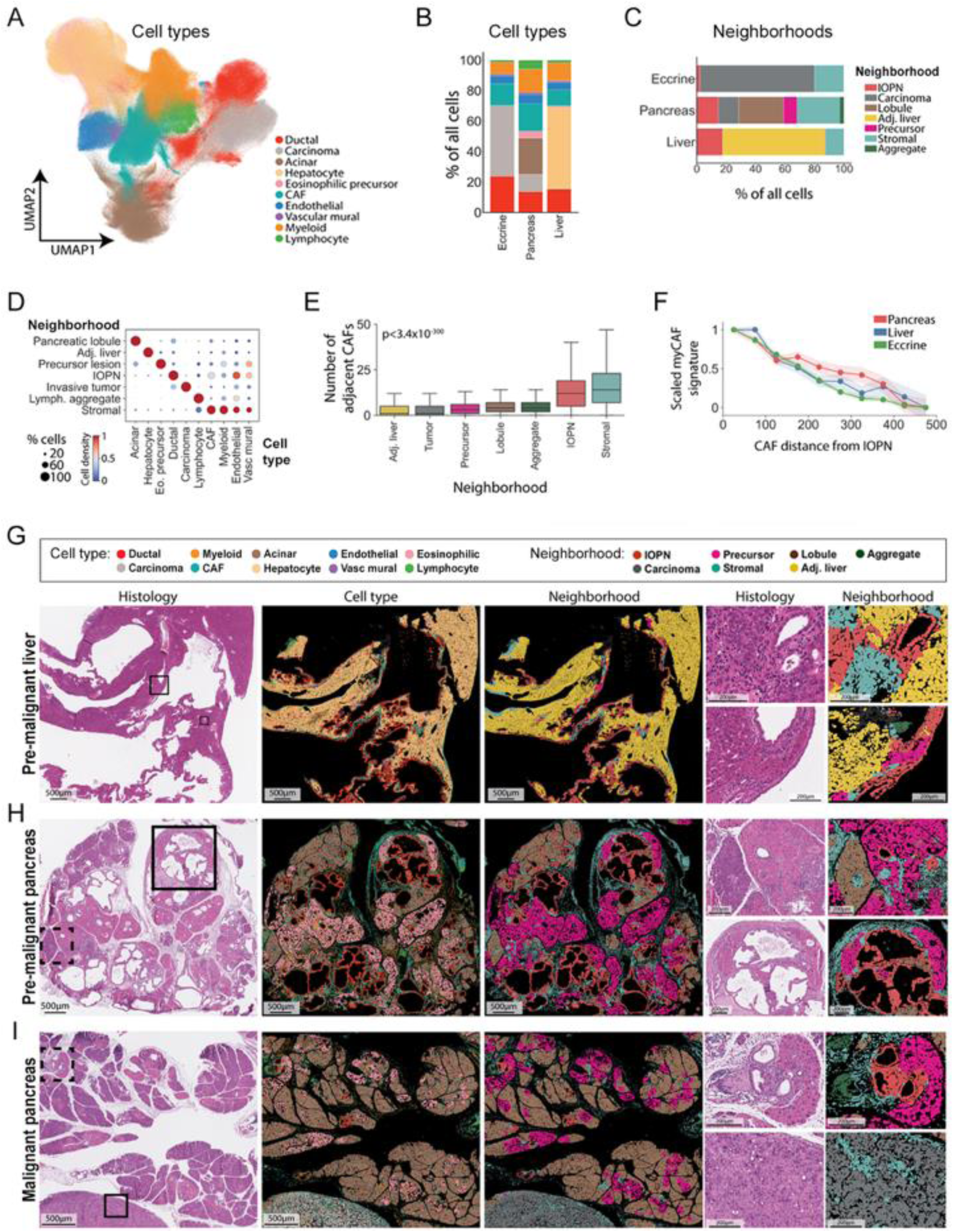
The evolving tumor and immune microenvironment in *DNAJB1-PRKACA* driven neoplasms. **(A)** Uniform Manifold Approximation and Projection (UMAP) projection of 1,390,407 individual cells representing 15 mouse samples colored by broad cell type. **(B)** Cell type frequencies from the spatial transcriptomics samples across aggregate sample type. (**C**) Cell neighborhood frequencies from the spatial transcriptomics samples across aggregate sample type. **(D)** Depicts the relative frequency of different cell types (x axis) and corresponding neighborhood (y axis). **(E)** Number of cancer associated fibroblasts (CAFs) within 50μm of cells within each neighborhood. *P*-value from Kruskal-Wallis test. **(F)** Scaled expression of myofibroblastic CAF (myCAF) marker genes within CAFs as a function of distance from nearest IOPN cell. **(G-I)** Representative histology, cell type, and neighborhood for pre-malignant liver, pre-malignant pancreas, and malignant pancreas samples. Magnification views of the liver and pancreas (black boxes) depict the complex distribution of cellular neighborhoods in different pathologies.

To further understand the microenvironment of these tumors, we next identified 7 distinct cellular neighborhoods according to the frequency of other cell types within 50 μm of each reference cell. These neighborhoods correlated with the major regions on histology, including normal pancreatic lobules, normal adjacent liver, eosinophilic precursor lesions, pre-malignant IOPNs, invasive tumor, lymphoid aggregates, and stroma (**Figure 6C, Supplemental Figure 5C-D**). Each neighborhood was evaluated with respect to the ten different cell types (e.g., acinar, hepatocyte, eosinophilic precursor, CAF, etc.). We noted that the stromal neighborhood was largely comprised of CAFs in proximity to myeloid cells and endothelial cells, with lymphocytes evident (**Figure 6D, Supplemental Figure 5E-F**). Notably, there were fewer lymphocytes in the pre-invasive lesions (precursor and IOPN) than the stromal neighborhood and a relative paucity of lymphocytes in the invasive carcinoma neighborhood. This is concordant with recent work depicting reduced T-cell infiltration and associated immune suppression in human tumors harboring *DNAJB1-PRKACA*,^18, 34, 35, 37^ and that the immunosuppressive stroma is coordinated by CAFs and tumor associated macrophages (TAMs) that sequester immune effector cells within the stroma and away from tumor cells.^37^ To understand the onset of fibrous stroma specifically, we calculated the density of CAFs immediately adjacent to (within 50 μm) cells within each of these neighborhoods. We found that the stromal and IOPN neighborhoods had the highest density of CAFs (**Figure 6E).** Furthermore, the expression of myofibroblastic markers within CAFs (myCAF signature) was directly related to their distance from the nearest IOPN neighborhood across all disease sites (**Figure 6F)**. Visual inspection further confirmed these findings, with the peak density of fibroblasts appearing around pre-malignant IOPNs in both liver and pancreas samples (**Figure 6G-H**). Within malignant samples, stroma was found predominately within stromal bands surrounding areas of confluent invasive tumor cells as seen in human *DNAJB1-PRKACA* driven cancers (**Figure 6I**).

Our histologic sections showed that the fibrous stroma was firmly established in the pre-invasive IOPNs (**Figures 4**–**5**); meanwhile, the stromal neighborhood analysis showed strong co-localization of CAFs and myeloid cells within the stroma (**Figure 6G-I**). Combined, this suggests the tumor permissive CAFs and TAMs are embedded within the stroma prior to malignant transformation. We therefore assessed the spatial orientation of *Acta2*+ CAFs (activated myofibroblastic CAFs)^39^ and *Cd163*+ TAMs (M2-like immunosuppressive TAMs)^40^ in pre-invasive liver IOPN and pancreas IOPN as compared to pancreas carcinoma. We examined three of our spatial sections including an advanced pre-malignant IOPN of the liver, (**Figure 7A**), advanced pre-malignant IOPN of the pancreas (**Figure 7B**) and invasive carcinoma (**Figure 7C**). Magnified views of the histologic sections demonstrated the complexity of each of these tumors including neoplastic cells / carcinoma cells and desmoplastic stroma (**Figure 7D-F**). Spatial analysis identified pre-malignant neoplastic cells in the liver IOPN and pancreas IOPN which matched the histologic identification of these cells (**Figure 7G-H**). Pre-malignant neoplastic cells were also evident in the periphery of the pancreas carcinoma section, but the bulk of the tumor consisted of malignant carcinoma cells (**Figure 7I**). Examination of the tumor microenvironment in these samples highlighted the proximity of the fibrotic stroma to the neoplastic cells (**Figure 7J-K**) and desmoplastic stroma to the carcinoma cells (**Figure 7L**). Consistent with our stromal neighborhood analysis, the stroma was comprised mostly of *Acta2*+ CAFs and *Cd163*+ TAMs, with T-cells confined to the stroma region. The proximity of these cell populations was likely coordinated by signaling molecules within the tumor microenvironment. The CAFs demonstrated high expression of signaling molecules such as *Fn1, Vegfc*, *Vegfd*, *Angptl4* and *Cxcl12* **(Supplemental Table 1**). *Cxcl12* is noteworthy due to the known interplay between CXCR4+ immune cells and the CXCL12+ CAFs in human tumors harboring *DNAJB1-*PRKACA.^37^ Furthermore, a prominent component of immune suppression in human *DNAJB1-PRKACA* driven cancer is due to immune exclusion orchestrated by *Cxcr4-Cxcl12* signaling in the tumor immune microenvironment.^37^ Indeed, there was robust expression of *Cxcr4* by the *Cd163*+ TAMs within the stroma in the pre-invasive liver IOPN (**Figure 7M**), pre-invasive pancreas IOPN (**Figure 7N**), and invasive carcinoma (**Figure 7O**). Taken together, these data support the idea that the developing fibrous stroma, comprised mostly of intercommunicating CAFs and TAMs, is a highly permissive environment that co-evolves with the *DNAJB1-PRKACA* expressing neoplastic cells (IOPN lesions), and likely facilitates invasive carcinoma development in this *DNAJB1-PRKACA* tumor model.

**Figure 7.**
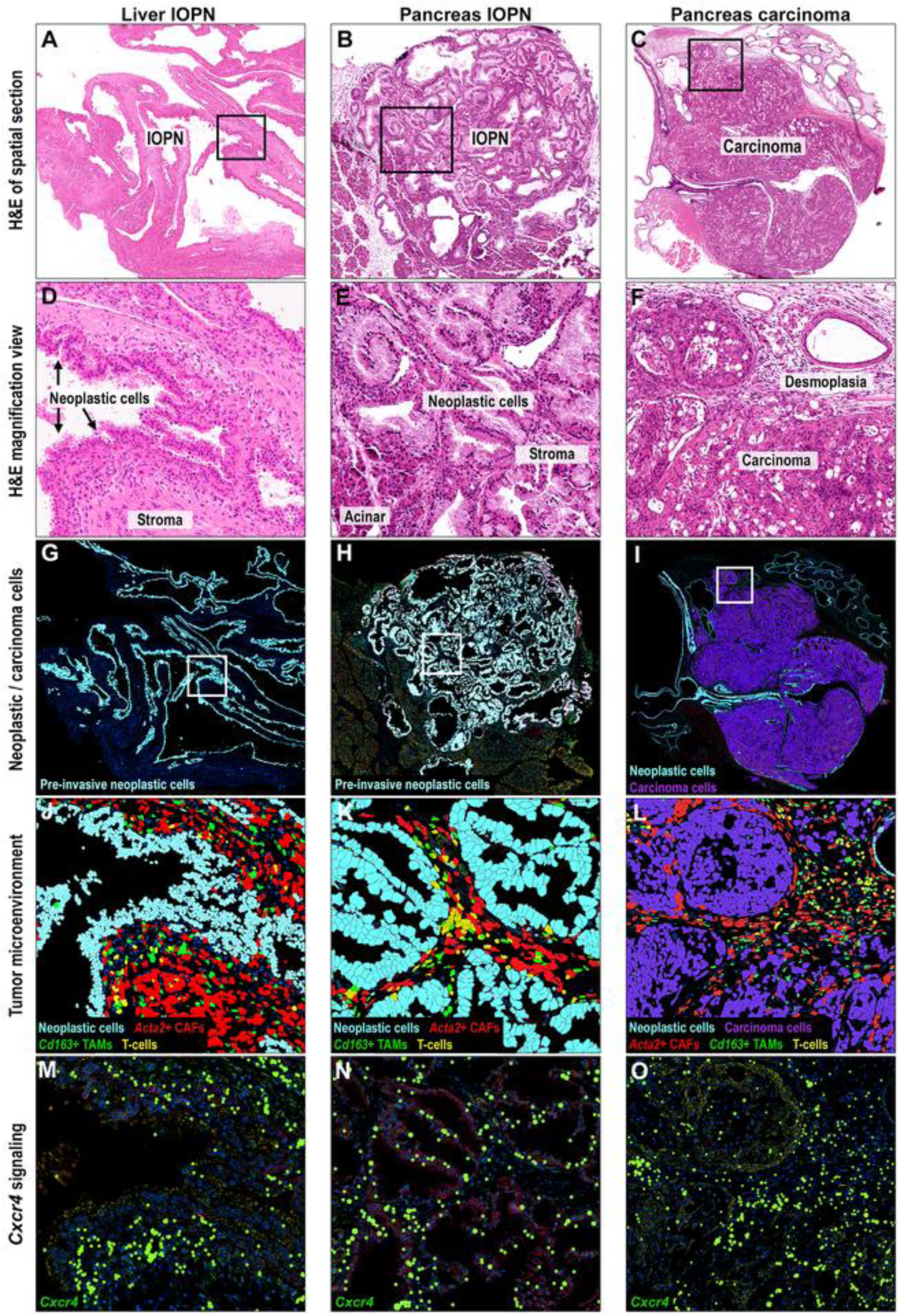
Spatial orientation of neoplastic and carcinoma cells in advanced IOPNs and carcinoma. Shown is H&E stain of spatial sections from an advanced IOPN of liver (**A**), advanced IOPN of pancreas (**B**), and pancreas invasive carcinoma (**C**) that formed from Ai14^D-P^; *Krt19^Cre:ERT^* mice approximately 4-5 months after tamoxifen administration. On H&E magnification views of the region in the black box, the architecture of the IOPN neoplastic cells adjacent to desmoplastic stroma is apparent in liver (**D**) and pancreas (**E**), with normal adjacent acinar cells captured in the pancreas section. (**F**) The pancreas carcinoma demonstrates dense desmoplasia surrounding the invasive carcinoma cells. (**G-I**) Spatial transcriptomics of the invasive carcinoma and surrounding region allows for highlighting of the pre-invasive neoplastic cells comprising the IOPNs as well as the carcinoma cells. (**J-L**) Both the IOPNs and the invasive cancer demonstrate significant infiltration of macrophages and fibroblasts in the tumor / immune microenvironment. Tumor associated macrophages (TAM, green) and cancer associate fibroblasts (CAF, red) are prominent cell populations within and surrounding neoplastic cells in pre-invasive lesions and in the stroma surrounding carcinoma cells. There is a relative paucity of T cells (yellow). **(M-O)** *Cxcr4* expression is observed in the stroma of pre-invasive IOPNs as well as pancreatic carcinoma.

### Molecular landscape in DNAJB1-PRKACA driven carcinoma uncovers an invasive biomarker

To characterize the molecular landscape among pre-invasive lesions and invasive carcinoma, several of our spatial transcriptomic samples made use of a custom panel to complement the standard spatial transcriptomics panel, including probes for *DNAJB1-PRKACA* and genes that are associated with human *DNAJB1-PRKACA* driven cancer (**Supplemental Table 2**).^41, 42^ This add-on panel was used for a subset of our spatial transcriptomic samples to generate n= 958,231 high-quality cells from 6 pancreatic and 6 liver samples (**Supplemental Figure 3A, 3C**) from tamoxifen-treated *Ai14^D-P^; Krt19^CreERT^* mice. We then directly compared invasive carcinoma and complex pre-invasive IOPN samples. We first evaluated the pancreas carcinoma (from **Figure 7**) for *DNAJB1-PRKACA* expression. As expected, there was robust expression of *DNAJB1-PRKACA* in the pancreas carcinoma (**Figure 8A**) and the distribution of *DNAJB1-PRKACA* expression signal precisely overlapped that of the carcinoma cells (**Figure 7I**). Human *DNAJB1-PRKACA*-driven carcinomas are strongly associated with several biomarkers caused by either the fusion oncoprotein expression in tumor cells or by the subverted stromal cells.^17, 32, 38, 41, 42^ For instance, *VDAC2* is firmly implicated in *DNAJB1-PRKACA*-induced metabolic dysfunction in human tumors,^17, 32^ and consequently, we identified robust expression of *Vdac2* in the cancer cells (**Figure 8B**). Meanwhile, *COL4A1* and *PDE10A* are *DNAJB1-PRKACA*-induced super enhancer associated genes that are consistently upregulated in human cancer and thought to be essential for cancer identity.^41, 42^ They are expressed in the stromal compartment (fibroblast) and by cancer cells, respectively,^41, 42^ which aligned with the pattern of *Col4a1* expression (**Figure 8C**) and *Pde10a* expression (**Figure 8D**) seen in our pancreas carcinoma.

**Figure 8.**
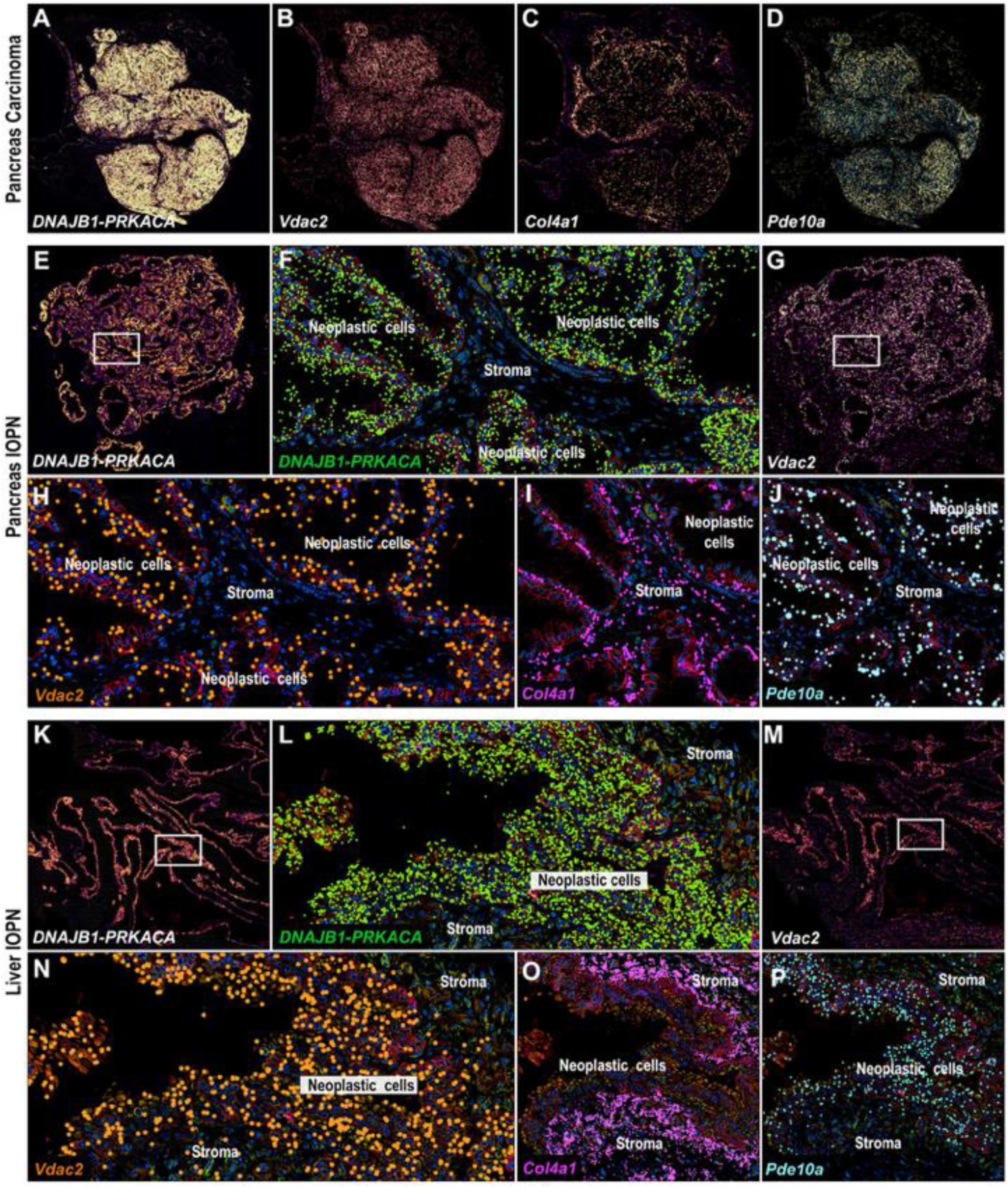
*DNAJB1-PRKACA* and associated gene expression in IOPNs and carcinoma. **(A)** *DNAJB1-PRKACA* is robustly expressed in pancreatic carcinoma cells. **(B)** *Vdac2* is observed to be highly expressed in carcinoma cells, suggesting metabolic dysfunction. **(C-D)** *Col4a1* and *Pde10a*, both super-enhancer associated genes consistently upregulated in human cancers. *Col4a1* expression overlaps with the stromal component while *Pde10a* is expressed in carcinoma cells, as shown. **(E-F)** *DNAJB1-PRKACA* expression is evident in the pancreas IOPN and corresponds to neoplastic cells. On higher magnification *DNAJB1-PRKACA* is notably absent in the stroma. **(G-H)** *Vdac2* expression correlates with the *DNAJB1-PRKACA* expressing cells in pancreatic IOPNs. **(I-J)** *Col4a1* expression corresponds to the stromal compartment while *Pde10a* expression corresponds to neoplastic cells. **(K-L)** *DNAJB1-PRKACA* expression is shown in liver IOPN cells and mimics the distribution seen in the pancreas IOPN, with localization to neoplastic cells. **(M-N)** *Vdac2* expression similarly is correlated with neoplastic cells in the liver IOPN. **(O-P)** Identical to the pancreatic IOPN, *Col4a1* expression corresponds to the stromal component while *Pde10a* expression corresponds to neoplastic cells.

In direct comparison, we evaluated the pre-invasive pancreas IOPN. The distribution of *DNAJB1-PRKACA* expression signal in the pancreas IOPN (**Figure 8E**) directly overlapped the neoplastic cells (**Figure 7H**), which mirrored the finding in the invasive pancreas carcinoma. Magnified views more clearly depicted the expression of *DNAJB1-PRKACA* in neoplastic cells and absence of expression in the stroma (**Figure 8F**, **Figure 7K**). Similarly, *Vdac2* expression was highly consistent with the distribution of *DNAJB1-PRKACA* expression (**Figure 8G**), which was confirmed on closer examination of the neoplastic cell / stroma interface (**Figure 8H**). Finally, *Col4a1* signal was robust within the stromal compartment between the nests of neoplastic cells (**Figure 8I**), while *Pde10a* expression was focused on the regions of neoplastic cells (**Figure 8J**). These findings were precisely reproduced in the pre- invasive liver IOPN, including *DNAJB1-PRKACA* distribution in neoplastic cells (**Figure 8K-L**, **Figure 7G**, **Figure 8J**), *Vdac2* expression strongly overlapping with *DNAJB1-PRKACA* expression (**Figure 8M-N**), and super enhancer associated genes *Col4a1* (**Figure 8O**) and *Pde10a* expression (**Figure 8P**) predominantly confined to the stroma and neoplastic cells, respectively. Consequently, we found that the pre-invasive IOPNs closely mimicked the invasive carcinoma with respect to many of the salient features associated with *DNAJB1-PRKACA*-driven cancer. This includes the oncocytic cellular phenotype, densely fibrotic stroma in the tumor microenvironment (e.g., CAFs, TAMs, collagen deposition), stromal gene expression (e.g., *Cxcr4*, *Col4a1*), and cancer cell autonomous gene expression (e.g., *Vdac2*, *Pde10a*). On further investigation, we did find one highly notable exception, which was *Slc16a14* expression.

*SLC16A14* is a known principal super-enhancer associated gene in human *DNAJB1-PRKACA*-driven carcinoma and uniformly upregulated regardless of the patient dataset.^22, 41, 42^ *SLC16A14* encodes for an orphan transporter in the monocarboxylase transporter family and its robust expression in human *DNAJB1-PRKACA*-driven cancer is intriguing due to the very low expression across normal tissues and other cancers.^22, 41^ Indeed, we found that *Slc16a14* was minimally expressed in normal cells (i.e., acinar cells and hepatocytes) with expression confined to the carcinoma cells (**Figure 9A**). Within the carcinoma cell population, there was a prominent sub-population of cancer cells that expressed high levels of proliferation markers (e.g., *Ki-67*, *Top2a*) suggestive of an aggressively proliferating cancer cell population (**Figure 9A**). Upon further examination, expression of *Slc16a14* in carcinoma cells was directly correlated with Ki-67 (*p* = 2.7×10^-168^ by Wald test, **Figure 9B**). To break this down further, we then reanalyzed three samples individually and reclassified cell types at more granular resolution. First, we used UMAP visualization of the various cell populations in the pancreas carcinoma sample to depict the carcinoma cells and highly proliferative carcinoma cells (**Figure 9C**) among the other cell populations within the tumor. Overlay of *DNAJB1-PRKACA* expression (**Figure 9D**) and *Slc16a14* expression on the same UMAP plot confirmed robust overlap of *Slc16a14* expression with the highly proliferative *DNAJB1-PRKACA* expressing carcinoma cells (**Figure 9E**). Using the same approach, we evaluated the pre-invasive pancreas IOPN and pre-invasive liver IOPN to depict the various cell populations (**Figure 9F**, **9I**) and *DNAJB1-PRKACA* expressing cells (**Figure 9G**, **9J**). In contrast to the invasive carcinoma, *Slc16a14* expression was essentially absent in the pre-malignant IOPNs, despite the presence of robust *DNAJB1-PRKACA* expression in the pre-invasive neoplastic cells (**Figure 9H**, **9K**). Thus, *Slc16a14* expression is restricted to the *DNAJB1-PRKACA* expressing cancer cells (cancer cell autonomous) and appears to signal the transition from pre-invasive tumor to invasive carcinoma (i.e. biomarker for invasion). These results substantiate that SLC16A14 is a highly promising target in *DNAJB1-PRKACA* driven cancer, which has been posited by other lead investigators in the field.^22, 41^ Taken together, we show how our novel *DNAJB1-PRKACA* model system can enhance our understanding of carcinogenesis (eosinophilic cells, desmoplastic stroma, invasive carcinoma), the tumor microenvironment (signaling), and the understanding of *DNAJB1-PRKACA* specific biomarkers (e.g., *Vdac2*, *Pde10a*, *Slc16a14*).

**Figure 9.**
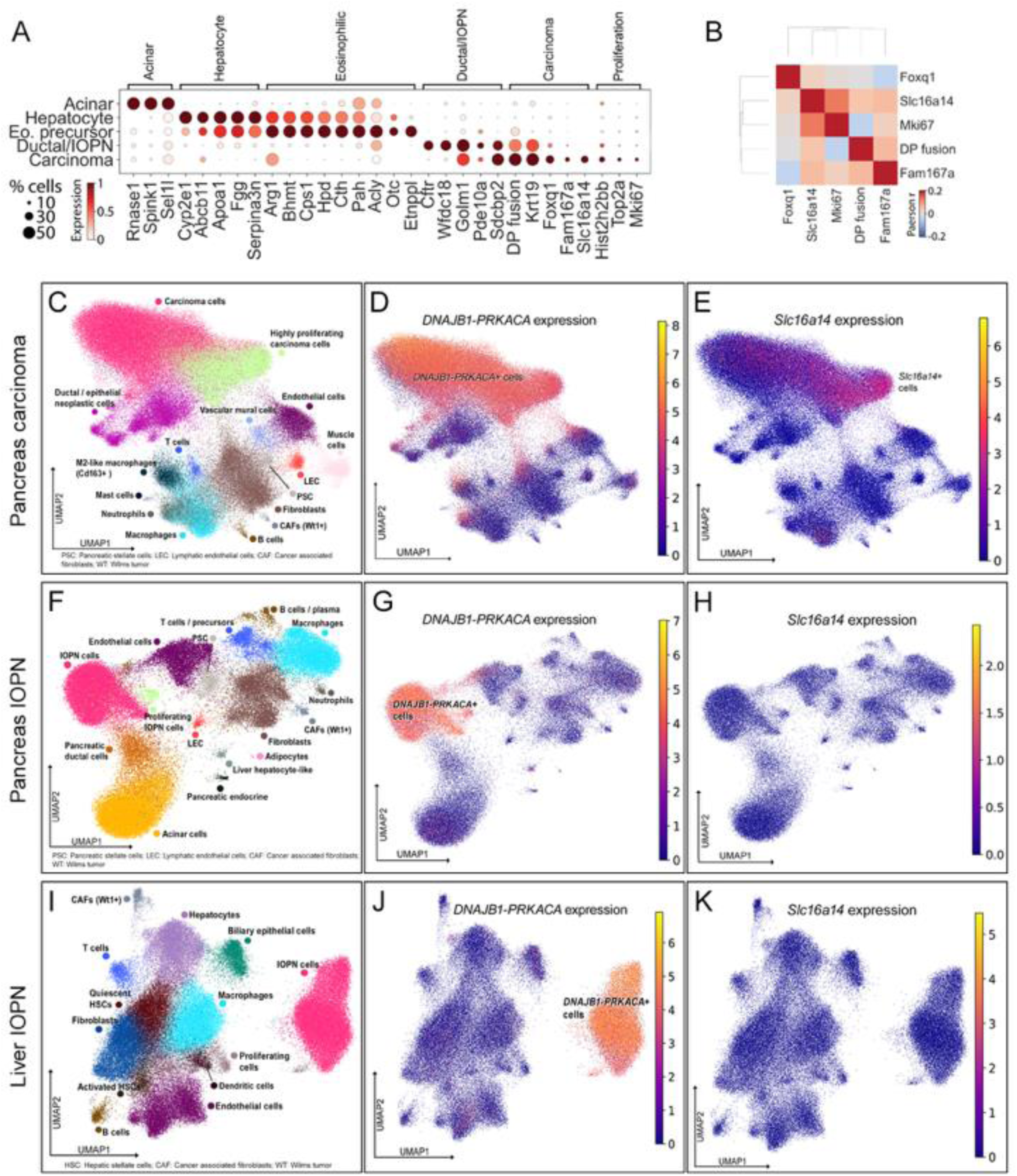
*Slc16a14* as a biomarker of invasive fusion-driven carcinoma. **(A)** Expression of selected marker genes for epithelial populations for spatial transcriptomics samples with custom add-on panel. **(B)** Significant Pearson correlations between expression of selected genes with carcinoma cells. (**C-K**). UMAP cell clustering of the spatial transcriptomics sample samples including pancreas carcinoma (**C**), pancreas IOPN (**F**) and liver IOPN (**I**). These pertain to the samples in Figures 7 and 8. Highlighted are the *DNAJB1-PRKACA* expressing cells (**D**, **G**, **J**) and *Slc16a14* expressing cells (**E**, **H**, **K**) in each sample. Expression levels of *DNAJB1-PRKACA* and *Slc16a14* are depicted by color scale which reveals robust expression of *DNAJB1-PRKACA* in each sample, but absence of *Slc1614* expression in the pre-invasive samples.

## Discussion

In a pair of studies from 2020,^1, 2^ the authors from each investigation identified that the *DNAJB1–PRKACA* gene fusion was present in intraductal oncocytic papillary neoplasms (IOPNs) of the pancreas and bile duct, and IOPN-associated cancer. Since then, additional studies have corroborated these results.^3, 4^ This compelling human correlative evidence uncovered a possible link between the *DNAJB1–PRKACA* gene fusion and the formation of subtypes of CCA and pancreatic carcinoma. Confirming that this *PRKACA* gene fusion causes rare subtypes of CCA and pancreatic carcinoma (IOPN-associated carcinomas) in an *in vivo* model system would therefore have profound implications for future clinical translation, particularly due to current precision medicine efforts to develop therapies targeting the fusion in FLC, another *DNAJB1-PRKACA*-driven cancer.^43, 44^ To investigate whether *DNAJB1–PRKACA* is a driver of IOPN and IOPN-associated carcinoma, we generated a genetically engineered mouse model allowing for cell-specific expression of human *DNAJB1– PRKACA* (conditional allele). The potential translatability of this approach is illustrated by other investigators who developed a conditional *in vivo* model to investigate another subtype of CCA, *IDH1*-mutant CCA.^45^ The constructed *in vivo* model allowed swift progress in understanding how the tumor cells orchestrated suppression of the immune system. This mechanistic insight could not be achieved in an immunosuppressed model, and evaluation of the tumor cells in an immune competent environment led to novel drug treatment combinations. Given the rare but deadly nature of FLC and IOPN-associated biliary and pancreatic carcinomas, a robust pre-clinical model is necessary to progress the understanding of disease biology and develop novel therapeutic strategies.^2, 46, 47^

Through our studies, we provide compelling evidence that the *DNAJB1–PRKACA* oncogene fusion alone drives a systematic and characteristic progression from eosinophilic neoplastic cell to IOPN to invasive carcinoma in murine liver (biliary tree) and pancreas (pancreatic duct). Our model demonstrates that the development of granular eosinophilic (oncocytic) cells with enlarged nuclei is the initial histologic manifestation of *DNAJB1-PRKACA* expression in both liver and pancreas, indicating that the metabolic (mitochondrial) and transcriptional changes are established at the earliest stage of tumor initiation. These oncocytic cells are a defining feature of human IOPN-associated carcinoma^3, 5, 9, 33^ and FLC^16^ and indicates an abundance of aberrantly functioning mitochondria in the cytoplasm.^48^ Electron microscopy in human IOPNs^3^, human FLC^3, 18, 49, 50^ and *DNAJB1-PRKACA* expressing tumors in mice^13^ confirm the presence of densely packed mitochondria with abnormal appearance (loss of cristae) and aberrant metabolism. Concordantly, mitochondrial dysfunction is a well-established hallmark of human FLC which has led to investigation of metabolic targets that can serve as a precision therapeutic strategy in *DNAJB1-PRKACA* driven cancers.^17, 18, 32, 51^ One such promising target is VDAC2 (voltage-dependent anion channel 2), a mitochondrial membrane channel protein that is uniformly increased in human FLC tumors (both at the transcript and protein level).^32^ VDAC2 dynamically changes sub-states to transport a variety of metabolic substrates and serves as a key regulator of mitochondrial pore permeability.^52^ A parallel critical function is to sequester BAK (BCL-2 homologous antagonist/killer) to regulate apoptosis.^53^ In turn, dysregulation of VDAC2 was associated with tumor cell metabolic dysfunction and survival in human FLC tumor slices.^17^ The connection between VDAC2 and metabolic dysfunction is notable because of the strong link between metabolic dysfunction and immunosuppression in human FLC.^17, 18^ Concordantly, *Vdac2* expression in tumor cells has been strongly linked to immune evasion and suppression of CD8+ T-cell immune effector response, indicating coordinate metabolic and immunosuppressive function in certain tumor types.^53^ In this manner, VDAC2 functions as an immune checkpoint, where inhibition enhances CD8+ T-cell mediated tumor cell death.^53^ Consistent with the human *DNAJB1-PRKACA* tumors, we found markedly increased *Vdac2* expression in our murine IOPN-associated carcinoma samples, and importantly, we also identified substantially increased *Vdac2* in the pre-cancerous neoplastic IOPN cells. Thus, the role of VDAC2 in *DNAJB1-PRKACA* driven cancer is likely established prior to carcinoma development, and future work using our model system will aim to delineate the putative link between metabolic dysregulation and immunosuppression.

In both liver (bile duct) and pancreas (pancreatic duct), we observed significant morphologic remodeling by *DNAJB1-PRKACA* expression. As IOPNs arose and became more advanced pre-cancerous lesions and eventually invasive carcinoma they developed a rich stroma characterized by significant desmoplasia. Desmoplastic stroma is found in human cancer including lamellar bands of dense fibrosis in FLC and hyalinized stroma in IOPN/IOPN-associated cancer.^2, 3, 7, 16^ How *DNAJB1-PRKACA* expressing tumor cells cause development of the desmoplastic stroma remains uncertain, but the connection between *DNAJB1-PRKACA* expressing cells, the desmoplastic environment and subsequent carcinoma is evident in our model system (i.e., carcinomas did not develop without fibrous stroma). In fact, this feature was also prominent in the *DNAJB1-PRKACA* induced soft tissue carcinomas we identified in our model, strongly supporting that the desmoplastic stroma is a hallmark feature of *DNAJB1-PRKACA* driven cancer in human patients and in mouse. Interestingly, prior mouse models of *DNAJB1-PRKACA* induced tumors showed eosinophilic (oncocytic) neoplasms that developed over a period of 14-20 months, but without a desmoplastic tumor stroma, which is an acknowledged limitation in these models.^13, 14, 18, 54^ We found that these eosinophilic neoplasms consisting of enlarged oncocytic cells were the initial changes that occurred (i.e., within 1-2 months) in both liver and pancreas due to *DNAJB1-PRKACA*. These early lesions progressed over time in our model to pre-cancerous IOPNs, which ultimately matured into carcinoma at ∼5 months after tamoxifen activation of *DNAJB1-PRKACA*. It is unclear why the prior model systems did not show a similar progression to densely fibrotic stroma or invasive carcinoma. One plausible explanation is cell of origin. While prior models used a hydrodynamic approach to introduce the oncogene fusion into liver cells,^13, 14^ we purposefully targeted the *DNAJB1-PRKACA* allele to biliary duct and pancreatic duct epithelial cells. From these cells emerged tumors that demonstrated strong biological overlap with human *DNAJB1-PRKACA*-driven cancer including cytologic features, transcriptional changes, and tumor stroma. Thus, our model is differentiated by *DNAJB1-PRKACA*-driven desmoplasia and carcinoma development, and we conclude that the desmoplastic stroma is of critical importance for driving the carcinoma phenotype.

The emphasis on a desmoplastic tumor microenvironment is due to its role in promoting tumorigenesis in a variety of cancers including cholangiocarcinoma^55–57^, pancreatic cancer^58, 59^ and FLC. Embedded within the desmoplasia of human *DNAJB1-PRKACA* driven cancers are an abundance of cancer associated fibroblasts (CAFs), which deposit type I, III and IV collagen,^7, 42^ and tumor associated macrophages (TAMs).^37^ For example, *COL1A1*, *COL1A2*, and *COL4A1* are highly expressed in human FLC,^42^ and these collagen types featured prominently in our murine tumors. Combined, CAFs and TAMs cooperatively foster an immune suppressed TME through coordinated immune effector cell dysfunction (exhaustion) and immune cell exclusion, which allows for cancer progression.^18, 34, 36, 37, 42^ For instance, T cell exclusion is promoted by stromal cells communicating via chemokine-receptor pairs, such as the chemokine CXCL12 and its receptor CXCR4.^37^ This chemokine ligand-receptor pair is upregulated in a variety of cancers and is correlated with increased metastatic spread and low overall survival.^60, 61^ Prior work in human FLC tumor slices has shown that CXCL12 production by CAFs in the desmoplastic stroma fosters sequestration of CXCR4 positive T cells to fibrotic areas of the tumor and away from tumor cells.^37^ Further, CXCL12 has been shown in other cancers to recruit CXCR4 positive tumor cells producing *VEGF* and CXCR4 positive TAMs producing *EGF*, resulting in endothelial cell recruitment and promotion of angiogenesis.^60^ Mirroring the human data, we observed that the TAMs expressed high levels of immunosuppressive markers (e.g., *Pdl1*, *Pdl2*, *Csf1r*), and that the TAMs / endothelial cells robustly expressed the chemokine receptor *Cxcr4* in proximity to the *Cxcl12*+ CAFs within the tumor and immune microenvironment. This data supports that the immunosuppressive microenvironment orchestrated by the stromal cells is achieved prior to the onset of invasive carcinoma. The CXCR4-CXCL12 axis is of particular interest because work investigating CXCR4 inhibition has already shown promise in potentiating immunotherapy in pancreatic cancer^62^ and FLC.^37^ Thus, the immune competent nature of our preclinical model will facilitate future work to examine the immune checkpoints that may be targeted in *DNAJB1-PRKACA* driven cancers.

Further, our spatial analysis enabled us to assess cell-autonomous expression of genes relevant to and consistently upregulated in *DNAJB1-PRKACA* driven cancer,^17, 32, 41, 63^ in both pre-cancerous tumors (IOPN) and invasive carcinoma. This is of critical clinical importance as *DNAJB1-PRKACA* driven cancers are genetically distinct from other more common carcinomas of the pancreas, liver and bile duct.^46, 64^ Outside of the *DNAJB1-PRKACA* gene fusion, there are no other recurring mutations identified (e.g., *KRAS*-mutation, *IDH1*-mutation).^8, 12, 13, 15^ Therefore, new therapies entering the clinic for other types of bile duct and pancreas cancer (e.g., mutant-KRAS inhibitors) are unlikely to prove effective in *DNAJB1-PRKACA*-driven carcinoma. Consequently, much work has been done to uncover promising downstream gene targets that appear critical for cancer cell identity (e.g., *DNAJB1-PRKACA* super enhancer associated genes), such that new therapies can be developed.^17, 19, 22, 32, 41, 42, 51, 63, 65, 66^ Examples include *COL1A1, COL4A1* and *VCAN* which are expressed by the CAF population, or *LINC00473*, *FAM167A*, *PDE10A, VDAC2, HK2,* and *SLC16A14*, which appear to be cancer-cell autonomous in our tumors. *LINC00473* was not identified, but this is because there is no murine homolog of this long non-coding RNA. Rather, recent evidence suggests that the ancestral *Pde10a* gene carried by mice split into *LINC00473* and *PDE10A* (Phosphodiesterase 10A) in humans.^67^ PDE10A is known to hydrolyze cyclic-AMP (cAMP) second messenger, and cAMP does not properly compartmentalize in human *DNAJB1-PRKACA*-driven cancer due to the aberrantly functioning PKA-fusion.^68^ Since *Pde10a* gene is cAMP inducible, improper cAMP sequestration causing reciprocal *Pde10a* expression is a plausible explanation for the early (pre-cancerous) elevation of *Pde10a* in our model.^67^ In fact, nearly all of the canonical metabolic and transcriptional changes due to *DNAJB1-PRKACA* appear early in tumor development, prior to the onset of invasive carcinoma. However, we identified one notable exception: *Slc16a14*. It is unclear why *Slc16a14* is only upregulated in the invasive carcinoma cells, despite robust expression of *DNAJB1-PRKACA* in the pre-cancerous tumor cells. This finding is particularly striking because *SLC16A14* shows the strongest association with *DNAJB1-PRKACA* induced super enhancers.^38, 41^ SLC16A14 is part of the solute carrier (SLC) superfamily and SLC16 monocarboxylate transporter (MCT) family of proteins, which include members that transport lactate, pyruvate and ketone bodies, among other key metabolic solutes.^69–74^ Little is known about SLC16A14, and there is a complete lack of knowledge about its function and transport substance, but what is known is of critical importance for SLC16A14 as a cancer target – there is a paucity of expression across most normal tissues.^69, 75^ Our findings of minimal *Slc16a14* expression in normal liver, pancreas and stromal cells align with the human data. In stark contrast, *SLC16A14* is markedly upregulated in *DNAJB1-PRKACA* driven tumors.^22, 38, 41, 42^ This suggests a remarkably unique biomarker with diagnostic and therapeutic implications – a biomarker that is firmly linked to *DNAJB1-PRKACA* expressing carcinoma cells (cancer cell autonomous) and is only overexpressed once invasive carcinoma forms. Commensurate with the role of the SLC superfamily as ‘metabolic gatekeepers of the cell’,^76^ SLC16A14 likely transports a substrate of critical value to the carcinoma cells, particularly given our finding of strong association with proliferative markers. Determining the transport substrate and function of SLC16A14 solute carrier protein in *DNAJB1-PRKACA* expressing carcinoma will be the focus of future research efforts.

Despite the compelling findings presented in this work, a key question that remains is why *DNAJB1-PRKACA* expression results in a variety of different clinical entities, including IOPN-associated cholangiocarcinoma, IOPN-associated pancreatic carcinoma and FLC. These findings suggest these diseases are phenocopies of one another and there may be additional *DNAJB1-PRKACA* driven carcinomas that are not well characterized. The data presented here support the clinical observation that *DNAJB1-PRKACA* is a bona fide driver of biliary and pancreatic IOPNs that can progress to carcinoma. Our mouse model faithfully recapitulates human disease through early metabolic dysfunction, the establishment of a desmoplastic stroma and immune suppressive microenvironment and expression of super-enhancer genes previously implicated in human *DNAJB1-PRKACA* driven cancers.

## Methods

### Animal Care

Animal studies were conducted at the University of Wisconsin in accordance with an approved animal protocol (M005959) by the University of Wisconsin School of Medicine and Public Health Institutional Animal Care and Use Committee. Mice were housed in an Association for Assessment and Accreditation of Laboratory Animal Care-accredited selective pathogen-free facility (UW Medical Sciences Center) on corncob bedding with chow diet (5020-Mouse Diet 9f; LabDiet, Richmond, IN) and water ad libitum. Ai14 tdTomato fluorescent reporter mice (*Ai14*, *B6.Cg-Gt(ROSA)26Sor^tm14(CAG-tdTomato)Hze^/J*) and Keratin 19-Cre^ERT^ knockin mice (*K19^CreERT^, Krt19^tm1(cre/ERT)Ggu^/J*) mice were purchased from the Jackson Laboratory (Bar Harbor, ME). All mice are congenic on a C57BL/6J background and were housed under identical conditions. The health and well-being of these animals was monitored closely by research and veterinary staff. Mice that showed signs of distress including weight loss (>20% total body weight), poor feeding, poor mobility, hunching or Body Condition Score less than 2 were euthanized via CO_2_ asphyxiation.

### Generation of the Rosa26^CAG-LSL-DNAJB1-PRKACA-IRES-tdTomato^ (Ai14^D-P^) mouse

The *Rosa26^CAG-LSL-DNAJB1-PRKACA-IRES-tdTomato^* (*Ai14^D-P^*) mouse was generated through the Advanced Genome Editing Core and Animal Models Core at the University of Wisconsin-Madison. To optimize human DNAJB1-PRKACA expression in the mouse, the codon-optimized cDNA was designed, synthesized and inserted into the pTARGATT6.1 vector (Applied StemCell, Milpitas, CA) by Sybio Technologies (Monmouth Junction, NJ). The internal ribosome entry site (IRES2) sequence was synthesized by Twist Bioscience (South San Francisco, CA) and inserted into the downstream of the *DNAJB1-PRKACA* cDNA sequence. The *Ai14^D-P^* allele was created by modifying the Ai14 allele by inserting a DNAJB1-PRKACA-IRES sequence, targeting upstream of the tdTomato coding sequence (Nterm.2: 5’-CTCCTCGCCCTTGCTCACCA-3’). This allele was created using a strategy adapted from the highly efficient gene insertion and editing approaches in mouse embryos.^77, 78^ Biotinylated donor was generated by amplifying the synthesized donor sequence using biotinylated primers **(Supplemental Table 3)**. *Ai14* mice were superovulated using Pregnant Mare Serum Gonadotropin (ProSpec Bio, East Brunswick, New Jersey) and human chorionic gonadotropin (Sigma-Aldrich, St. Louis, Missouri), and then mated. Day 0.5 embryos were injected with 10 ng/μl guide RNA, 20 ng/μl biotinylated donor, and 37.5 ng/μl Cas9 mRNA (Aldevron, Madison, WI). Injected embryos were transplanted into pseudopregnant B6D2F1 females and pups were sequenced at weaning to assess for gene edits.

To characterize genome editing events, a short fragment of the knockin cassette was amplified (primers 2381/2382) **(Supplemental Table 3)**. Samples carrying the small fragment were amplified with longer range PCR targeting the 5’ (primers 1968/2382) and 3’ (primers 2381/1969) ends **(Supplemental Table 3)**, with one primer located within the knockin cassette and the other primer outside of the homology arms, and samples yielding bands of expected size were Sanger sequenced (DNA Sequencing Facility). F0 animals carrying the expected sequence were backcrossed to C57BL/6J mice. F1s were sampled and 5’ and 3’ PCRs were performed as above. Amplicons of expected size were fragmented (DNA Fragmentation Kit (6137),Takara), end repaired (NEBNext Ultra II End Repair/dA-Tailing Module, NEB), ligated (NEBNext Ultra II Ligation Module (E7546), NEB) to adapters (xGen Stubby Adapter (E7595), IDT), libraries were amplified (primers 844/845) **(Supplemental Table 3)** and indexed with index PCR, and sequenced on a MiSeq 2 x 250 Nano, and assembled with SPAdes de novo assembler.^79^ During this characterization, it was revealed three base pairs (corresponding to Val2) were missing from the tdTomato coding sequence.

### Genomic characterization of Ai14^D-P^ mice

To validate the Ai14^D-P^ allele against large-scale rearrangements, duplications, and inversions arising from double-strand break repair that are undetectable by PCR^28, 29, 80–82^, the focal F1 was sequenced by whole-genome long-read Oxford Nanopore Technology (ONT), yielding 11.3X read depth at chromosome 6 after alignment to GRCm39 (Advanced Genome Editing Laboratory). A 100 kbp window centered on the Cas9 cleavage site was assessed for structural variants using cuteSV and manually inspected for soft-clipped reads indicative of unexpected genomic insertions. Predicted off-target sites were identified by intersecting CRISPOR^83^ and Cas-OFFinder^84^ outputs, retaining loci with ≤2 mismatches and ≤1 RNA/DNA bulges. No indels were observed at any predicted off-target loci. To confirm the precise arrangement of the *Ai14^D-P^* allele, reads intersection Gt(ROSA)26Sor (GRCm39:6:113043656-113059075), were subsetted and aligned to the expected sequence **(Supplemental Table 3)** with minimap2. This analysis revealed a duplication of the knock-in sequence that was undetectable by PCR above. The resultant allele carried two DNAJB1-PRKACA fusion coding sequences and a tdTomato coding sequence, each separated by a polycistronic IRES cassette. Five contiguous reads spanning the entire donor construct confirmed the duplication.

To confirm the Loxp-Stop-LoxP and DNAJB1-PRKACA-IRES-tdTomato cassettes in *Ai14^D-P^* allele, PCR was performed with forward primer (5’-LSL): 5’- ACGTGCTGGTTATTGTGCTGT-3’ and reverse primer (3’-tdTomato): 5’-TTACTTGTACAGCTCGTCCATGCC-3’. PCR was carried out for 30 cycles (98 °C for 10’ (1 cycle), 98 °C for 10’’, 60 °C for 15”, and 68 °C for 5’ (30 cycles)) using PrimeStar GXL DNA polymerase (Takara Bio, Kusatsu, Japan). The 6.4 kb amplicons were recovered from 1% agarose gel using QIAquick gel extraction kit (Qiagen, Venlo, Netherlands) after electrophoresis. The amplicons were digested with EcoRI (Promega, Madison, WI), and DNA fragments of LoxP-Stop-LoxP (1.1 kb), DNAJB1-PRKACA-IRES (2.0 kb), and DNAJB1-PRKACA-IRES-tdTomato (3.3 kb) were detected. To confirm excision of the stop cassette in the genome, the same PCR protocol was employed.

To determine the genotype of wild-type and *Ai14^D-P^* alleles, the standard genotyping PCR protocol for *Ai14* mice (Protocol 29436, The Jackson Laboratory) was used. The codon-optimized hDNAJB1-PRKACA cDNA construct was detected by PCR using forward primer (FLC3): 5’-CAGAAAGAGGGAGATCTTCGACCGC -3’ and reverse primer (FLC4): 5’- ACGTACTCCATGACCATGTACAGG-3’. PCR was carried out for 29 cycles (94 °C for 2’ (1 cycle), 94 °C for 30’’, 61 °C for 30”, and 72 °C for 30’ (29 cycles)) using GoTaq DNA polymerase (Promega). A 362-bp PCR product was amplified from the *Ai14^D-P^* alleles.

### Tamoxifen Induction

The *Ai14^D-P^* mice were crossed to *K19^CreERT^* mice to generate *Ai14^D-P^; K19^CreERT^* mice. Experimental animals were homozygous for *Ai14^D-P^* and heterozygous for *K19^CreERT^.* Tamoxifen stock solution was prepared using corn oil as a vehicle at a concentration of 20 mg/mL and 70 μL (55-70 mg/kg body weight) was administered every other day for a total of 5 doses via intraperitoneal injection in 6-8 week-old mice. Tissues were collected at monthly endpoints or once humane endpoints met for longer term studies.

### Histopathology, Immunohistochemistry and Immunofluorescence

The stomach, proximal small intestine, liver and pancreas were collected for histology at endpoint and placed in 10% Neutral Buffered Formalin (NBF) for 24 hours, then stored in 1X Phosphate-Buffered Saline (PBS) and kept at 4 °C. Tissues were then processed into paraffin on a Sakura VIP5 tissue processor. The tissue blocks were sectioned on a Leica RM2125RT microtome at section thickness of 5um and mounted on StatLab Colorview Adhesion slides. H&E and Immunohistochemistry were performed by UW-Translational Research Initiatives in Pathology Core facility (TRIP). For Immunohistochemistry, RFP (600-401-379, Rockland Immunochemicals, Pottstown, PA), Pan Cytokeratin (ab234297, Abcam, Cambrige, MA), and Ki-67 (D3B5) (#12202, Call Signaling Technology, Danvers, MA) antibodies were employed.

For preparation of Xenium slides, regions of interest on six different sample blocks were surface-scored using a 6 mm biopsy tool. Sections were cut at 5 μm and subsequent 6 mm circle sections were arranged in a 3 x 2 pattern on the Xenium slide. Slides were dried overnight in a dessicator. H&E stain was performed on the Xenium slide through a Leica Autostainer XL after the Xenium assay.

### Cre-expression using Adeno-associated Virus

CMV-promoter-controlled Cre recombinase expression (pENN.AAV.CMVs.Pl.Cre.rBG (#105537)) and null (pAAV.TBG.PI.Null.bGH (#105536)) adeno-associated viruses packaged in the capsid of serotype 8 (AAV8) were obtained from Addgene (Watertown, MA). The AAV8 particles (5 x 10^10^ genomic copies (GC)) were suspended in 100 μl of PBS(-) solution and injected into 9-week-old *Ai14^D-P^* mice through the retro-orbital sinus. Two weeks after the AAV injection, livers were collected. For immunoblotting, tissues were lysed and homogenized in RIPA lysis reagent (Thermo Fisher, Waltham, MA) with Halt protease and phosphatase inhibitor cocktail (Thermo Fisher). For fluorescence imaging, tissues were fixed in10% NBF for 24 hours and then stored in 30% sucrose at 4 °C. Fixed tissues were embedded in Tissue-Tek O.C.T compound (Sakura Finetek, Torrance, CA) and stored at -80 °C. Frozen tissues were sectioned at a section thickness of 10 μm, and tissue slides were prepared by the UW-Experimental Animal Pathology Laboratory. Images of red tdTomato-expressing cells were acquired using an A1R MP+ Confocal Microscope (Nikon, Shinagawa, Japan) with 561 nm laser. Images were analyzed using NIS-Elements C-ER software (Nikon).

### Sorting tdTomato-positive Cells

Four months after the tamoxifen injections in *Ai14^D-P^; K19^CreERT^* mice, liver were collected. Livers were minced into pieces 1 mm in diameter or less, rinsed in PBS and incubated at 37 °C in a solution of Collagenase P (Millipore Sigma, Burlington, MA) in HBSS at a concentration of 1 mg/ml for 30 minutes. The cells were then washed with R10 media and strained sequentially through a 500 μM filter, 100 μM filter and 40 μM filter to prepare a solution of dissociated cells. Dissociated liver cells were cultured in William’s E medium (Life Technologies, Carlsbad, CA) supplemented with 10% fetal bovine serum (Cytiva, Chicago, IL) and 1% penicillin–streptomycin-glutamine (Life Technologies) on Collagen I (Rat tail) -coated culture dishes (Corning, New York, NY) at 37 °C and 5% CO_2_. Six weeks after the cell culture, red fluorescent protein (tdTomato) positive and negative cells were sorted at 4°C using a BD FACSAria III (BD Biosciences, Franklin Lakes, NJ). For immunoblotting, cells were lysed and homogenized in RIPA lysis reagent (Thermo Fisher) with Halt protease and phosphatase inhibitor cocktail (Thermo Fisher). For genomic PCR, genomic DNA was isolated using GeneJet Genomic DNA purification kit (Thermo Fisher). Images of red tdTomato-expressing cells were acquired using an A1R MP+ Confocal Microscope (Nikon, Shinagawa, Japan) with 561 nm laser. Images were analyzed using NIS-Elements C-ER software (Nikon).

### Immunoblotting

Forty μg of liver or cell homogenate was loaded on Mini-Protean TGX protein gel (7.5%) (Bio-Rad, Hercules, CA) and transferred to Immobilon-P membranes (MilliporeSigma, Burlington, MA) by electrophoresis. Membranes were incubated in 10% w/v BSA blocking buffer (Thermo Fisher) at room temperature for 1.5 hour and hybridized with primary antibody (PKA(c): 1:2000, (610981, BD Biosciences, Franklin Lakes, NJ), β-Actin: 1:2000 (#4967, Cell Signaling Technology, Danvers, MA) at 4°C overnight. After hybridization with Alkaline Phosphatase (AP)-conjugated secondary antibody (anti-mouse: 1:5000 (#7056, Cell Signaling Technology, Danvers, MA)), anti-rabbit: 1:5000 (111-055-144, Jackson ImmunoResearch, West Grove, PA)), proteins were detected using NBT/BCIP solution (Thermo Fisher). The samples were loaded separately and membranes were developed independently. The images were analyzed using ImageJ software (Fiji).

### Spatial Transcriptomics

Spatial transcriptomics was performed using a custom panel built off the 10x Genomics Xenium Prime 5K Mouse Pan Tissue & Pathways Panel (10x Genomics, Pleasanton, CA). Gene panel information, including probe sequences and accession numbers, is provided in Supplementary Table 1. For preparation of Xenium slides, regions of interest on six different sample blocks were surface-scored using a 6mm biopsy tool. Sections were cut at 5 μm and subsequent 6mm circle sections were arranged in a 3 x 2 pattern on the Xenium slide. Slides were dried overnight in a dessicator. Xenium in situ analysis was performed using the 10x Genomics Xenium Analyzer instrument according to manufacturer recommendations. H&E stain was performed on the Xenium slide using a Leica Autostainer XL after the Xenium assay.

### Data Analysis

Cell segmentation was performed using the Xenium onboard analysis tool (v3.3) according to staining performed with the Multimodal In Situ Cell Segmentation Kit (10x Genomics, Pleasanton, CA). Gene expression for each segmented cell was exported for downstream bioinformatic analysis. All analysis was conducted using custom Python scripts and the Scanpy ecosystem.^85^ Stringent quality control was performed to remove segmented cells with inadequate cell area, total transcript count, or number of unique genes. Following normalization and log-transformation of transcript counts, batch correction was performed using HarmonyPy.^86^ Manual graph-based cell type annotation was performed using standard marker genes. Cell type abundances were compared with a Bayesian model for compositional single-cell analysis using scCODA.^87^ Cellular neighborhoods were identified by counting the number of cells within a 50μm radius of each reference cell. The resulting adjacency matrix was embedded using principal component analysis (PCA) and neighborhoods identified using a coarse-grained k-nearest neighbors clustering algorithm. The distance to the nearest neoplastic cell was calculated using the Scipy KDTree implementation.^88^ Distances were binned and a myCAF score calculated using the Scanpy score_genes function with the following gene set: *Col12a1, Col1a1, Col1a2, Col4a1, Col4a2, Col5a1, Fap, Lrrc15, Pdgfra, Plaur, Postn, Tagln,* and *Thbs1*. Data visualization was performed using the Xenium Explorer platform (10x Genomics, version 4.1.1).

Individual sample analyses in Figure 9 were conducted using Scanpy and spatialdata for Python.^85, 89^ Briefly, cell segmentations from default Xenium analysis were imported as AnnData objects into Scanpy.^90^ Cells were filtered based on a maximum 98th percentile cutoff of total counts and a minimum cutoff of 20 counts/cell. Genes detected in less than 100 total cells were also removed. Total counts per cell were normalized, log transformed and scaled. Nearest neighbors graph construction was performed using a cosine distance metric directly on the scaled expression matrix and UMAP was performed, followed by clustering with the leiden algorithm. Cells were annotated via a combination of automated prediction by CellTypist and manual marker gene evaluation.^91^ Where appropriate, subclustering analysis was performed by isolating individual cell populations and applying highly variable gene selection and manual inclusion of relevant marker genes, followed by PCA, neighbor graph construction, UMAP and reclustering with the leiden algorithm.^92, 93^

All raw and processed spatial transcriptomics data has been deposited to the NCBI Gene Expression Omnibus (GEO) at accession GSE326148. Raw sequencing data from the Ai14^D-P^ mouse model is available at NCBI Sequence Read Archive (SRA) at accession SRR37846526.

## Acknowledgement

This work was supported by the U.S. Department of Defense (RA210165, SMRK) (CA180067 VGP) (CA230890 JAC), Fibrolamellar Cancer Foundation (SMRK, VGP), and National Institutes of Health (T32CA090217, PRC, ACV).

The authors would like to acknowledge the University of Wisconsin School of Medicine and Public Health Biomedical Research Model Services and Translational Research in Pathology (TRIP) core facilities. The authors also acknowledge the University of Wisconsin-Madison Biotechnology Center Animal Models Core and Advanced Genome Editing Laboratory, Gene Expression Center and Bioinformatics Research Core. Finally, the authors thank Dr. Darya Buehler for her analysis of the soft tissue tumors.

## List of Abbreviations

AAV8: Adenovirus Associated Vector 8
CAF: Cancer Associated Fibroblast
CCA: Cholangiocarcinoma
FLC: Fibrolamellar Carcinoma
H&E: Hematoxylin & Eosin
IHC: Immunohistochemistry
IOPN: Intraductal Oncocytic Papillary Neoplasm
IP: Intraperitoneal
Krt19: Cytokeratin 19
PKA: Protein Kinase A
SLC: Solute Carrier
TAM: Tumor Associated Macrophage
TME: Tumor Microenvironment

---

**Supplemental Figure 1.**
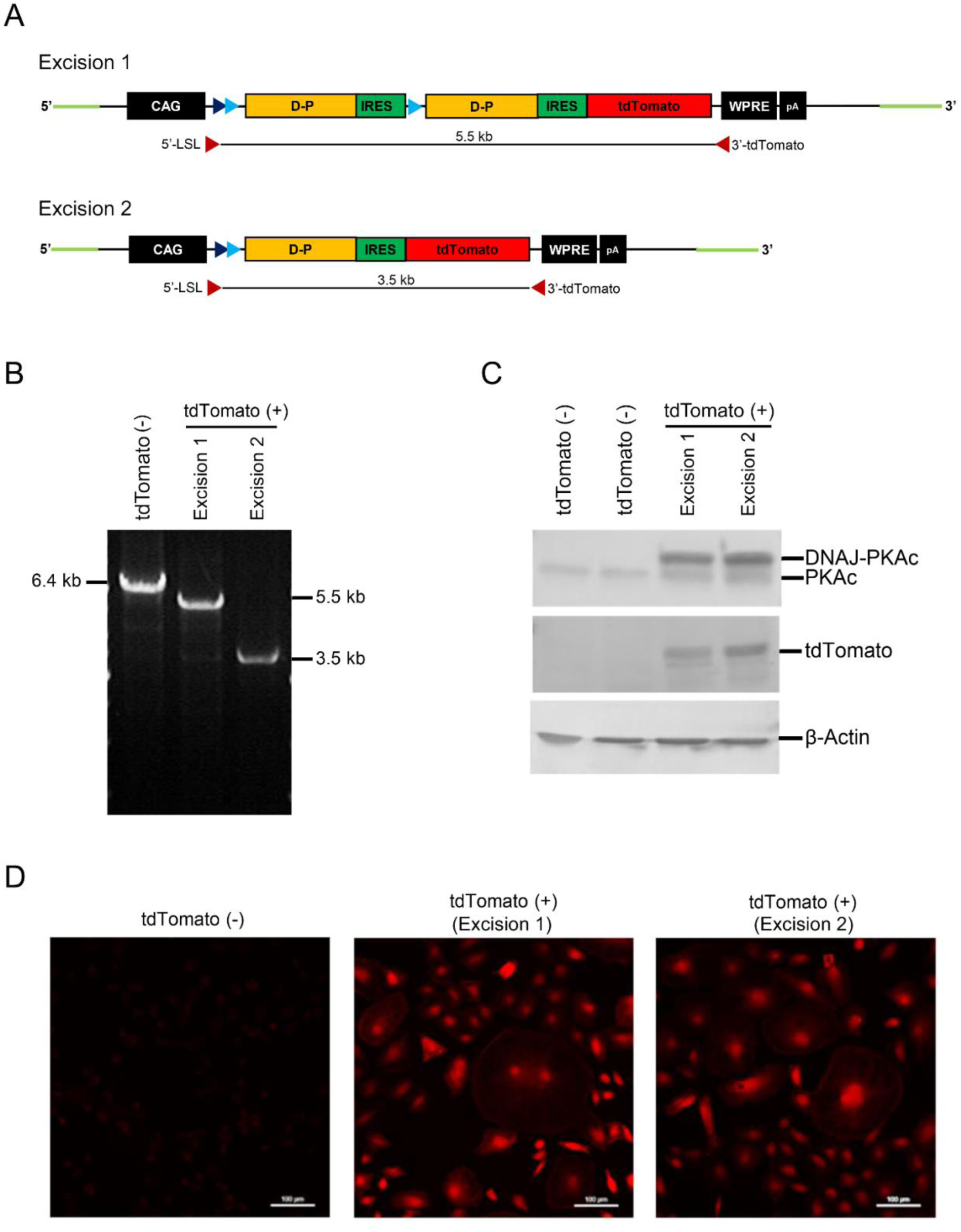
Activation of human DNAJB1-PRKACA expression-fluorescent reporter cassette by Cre-loxP recombination. **(A)** The hDNAJB1-PRKACA and tdTomato-positive cells show the excision of the Stop cassette (Ex-1) or the Stop-D-P-IRES cassette (Ex-2) flanked by loxP sites through Cre-loxP recombination. **(B-C)** The tdTomato-positive and negative cells were isolated from liver and pancreas of tamoxifen-treated *Ai14^D-P^; K19^CreERT^* mouse. **(B)** PCR screening, The 5.5 kb (Ex-1) and 3.5 kb (Ex-2) amplified bands were detected in tdTomato-positive cells. **(C)** Analysis for hDNAJB1-PRKACA expression, D-P proteins were detected using whole cell homogenate (20 μg) from tdTomato-negative (tdTomato (-)) and positive (tdTomato (+)) cells. The native PKAc and D-P protein was detected using PKAc antibody. β-actin was used as a housekeeping control. **(D)** The tdTomato-positive cells (red) were detected using a confocal microscope. Scale bar: 100 μm

**Supplemental Figure 2.**
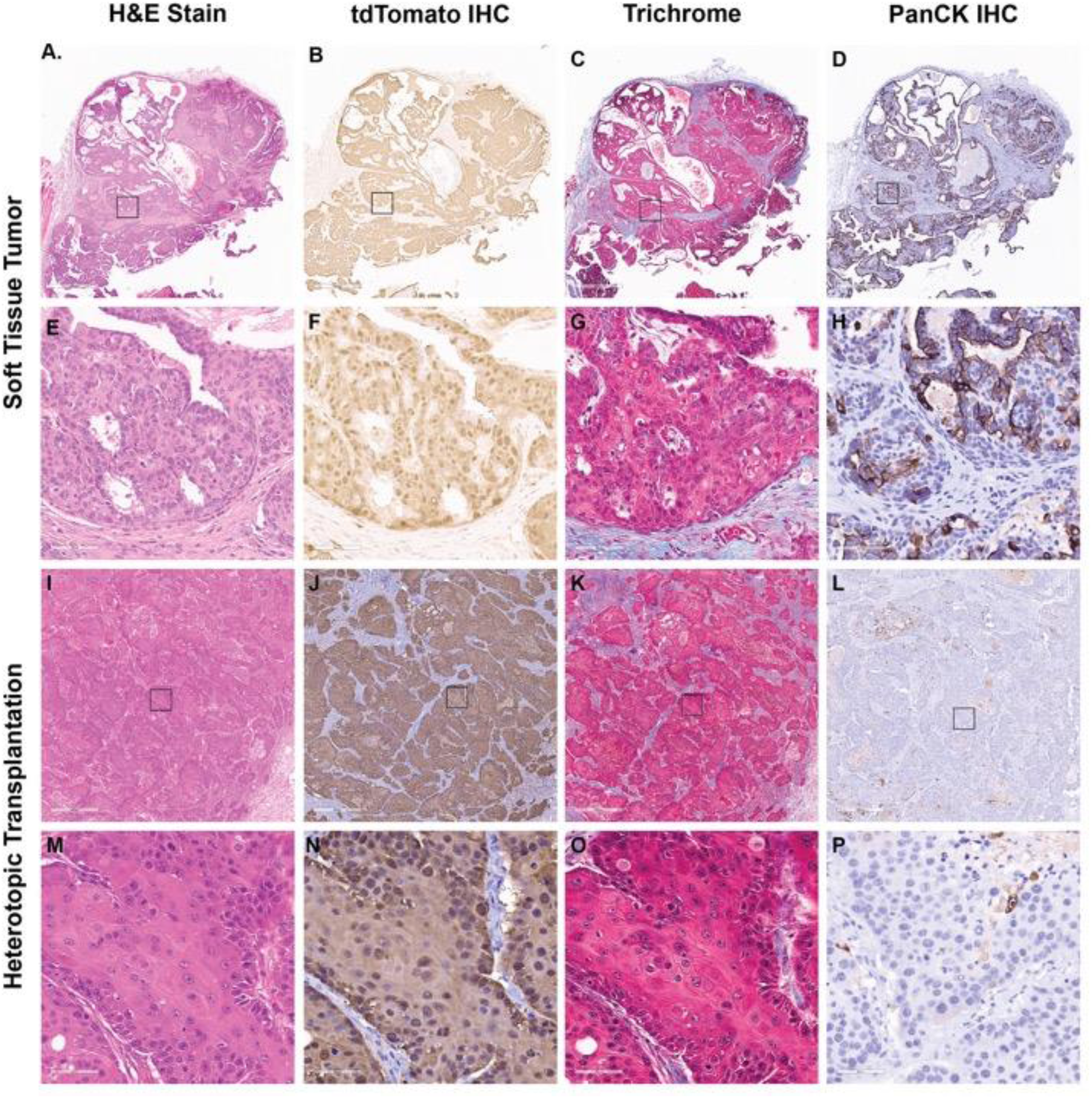
D*N*AJB1*-PRKACA*-driven soft tissue tumors. The primary soft tissue tumor isolated from the limb of a Ai14^D-P^; Krt19^Cre:ERT^ mouse is shown with subsequent passage of tumor in a C57Bl6 mouse. **(A)** H&E of primary soft tissue tumor with cystic regions similar to cystic lesions in pancreas and liver. **(B)** Immunohistochemical staining positive for tdTomato confirms expression of *DNAJB1-PRKACA*, **(C)** trichrome shows collagenous bands interspersed in tumor and **(D)** immunohistochemical stain for pancytokeratin confirms epithelial origin. **(E-H)** Magnification views of regions of interest in black boxes from panels **(A-D)**. **(I)** H&E of tumor from panel **(A-D)** following multiple heterotopic flank passages in a C57Bl6 mouse. The solid components of tumor predominate and there are fewer cystic regions than before. **(J)** Immunohistochemical staining positive for TdTomato confirms these cells retain *DNAJB1-PRKACA* expression, **(K)** trichrome shows persistence of collagenous bands in tumor and **(L)** immunohistochemistry for pancytokeratin is scant following subsequent passages but present. **(M-P)** Magnification views of regions of interest in balck boxes from panels **(I-L)**.

**Supplemental Figure 3.**
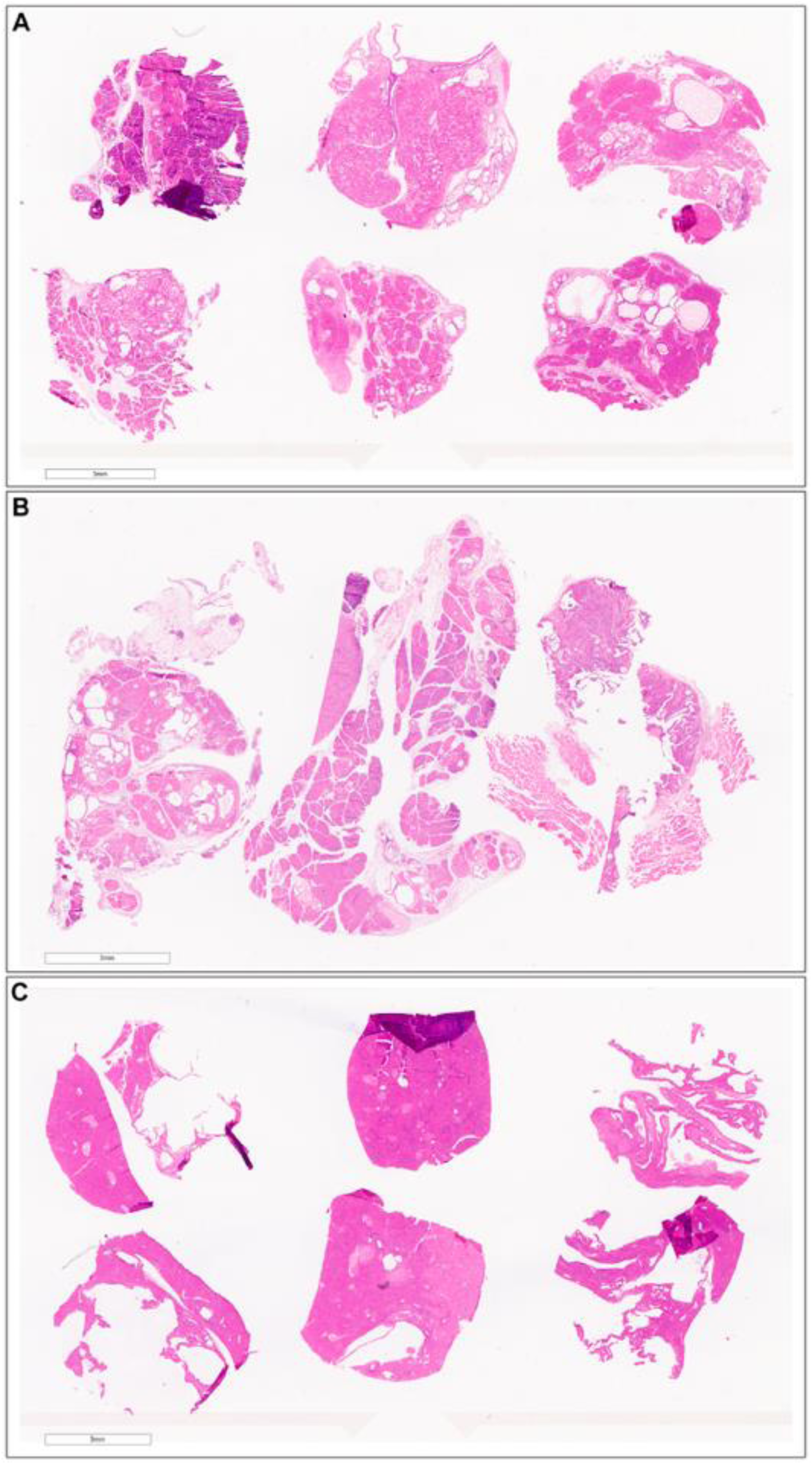
Spatial transcriptomics sections. Regions of interest in pathology sections from Ai14^D-P^; Krt19^Cre:ERT^ mice were identified, mounted to a Xenium slide, and spatial transcriptomics performed to evaluate different cell populations and RNA expression in the tumor / immune microenvironment. Shown is the baseline H&E of the sections for spatial transcriptomics including pancreas (**A**), pancreas and soft tissue (**B**), and liver (**C**). A custom Xenium panel including probes for *DNAJB1-PRKACA* was used for the pancreas (n = 6) and liver (n = 6) sections from **A** and **C**.

**Supplemental Figure 4.**
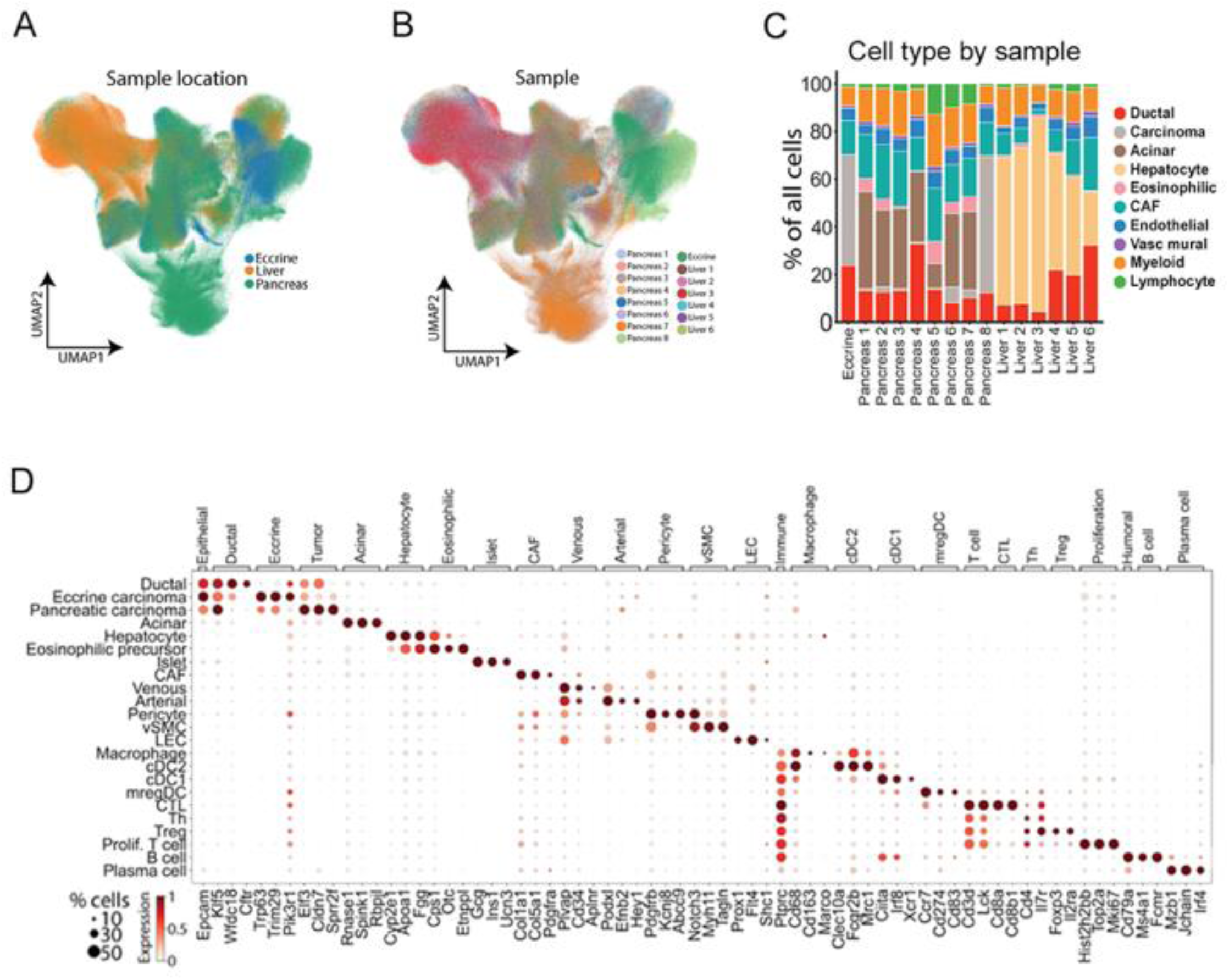
The evolving tumor and immune microenvironment in *DNAJB1-PRKACA* driven neoplasms. **(A)** Uniform Manifold Approximation and Projection (UMAP) projection of 1,390,407 individual cells representing 15 mouse samples colored by broad cell type. **(B-C)** Cell type frequencies from the spatial transcriptomics samples across sample type **(B)** and between individual samples **(C)**. **(D)** Expression of selected marker genes for each cell type with marker genes on the x-axis and cell type on the y-axis. Scale representing percent of cells expressing the marker gene and expression level is also depicted.

**Supplemental Figure 5.**
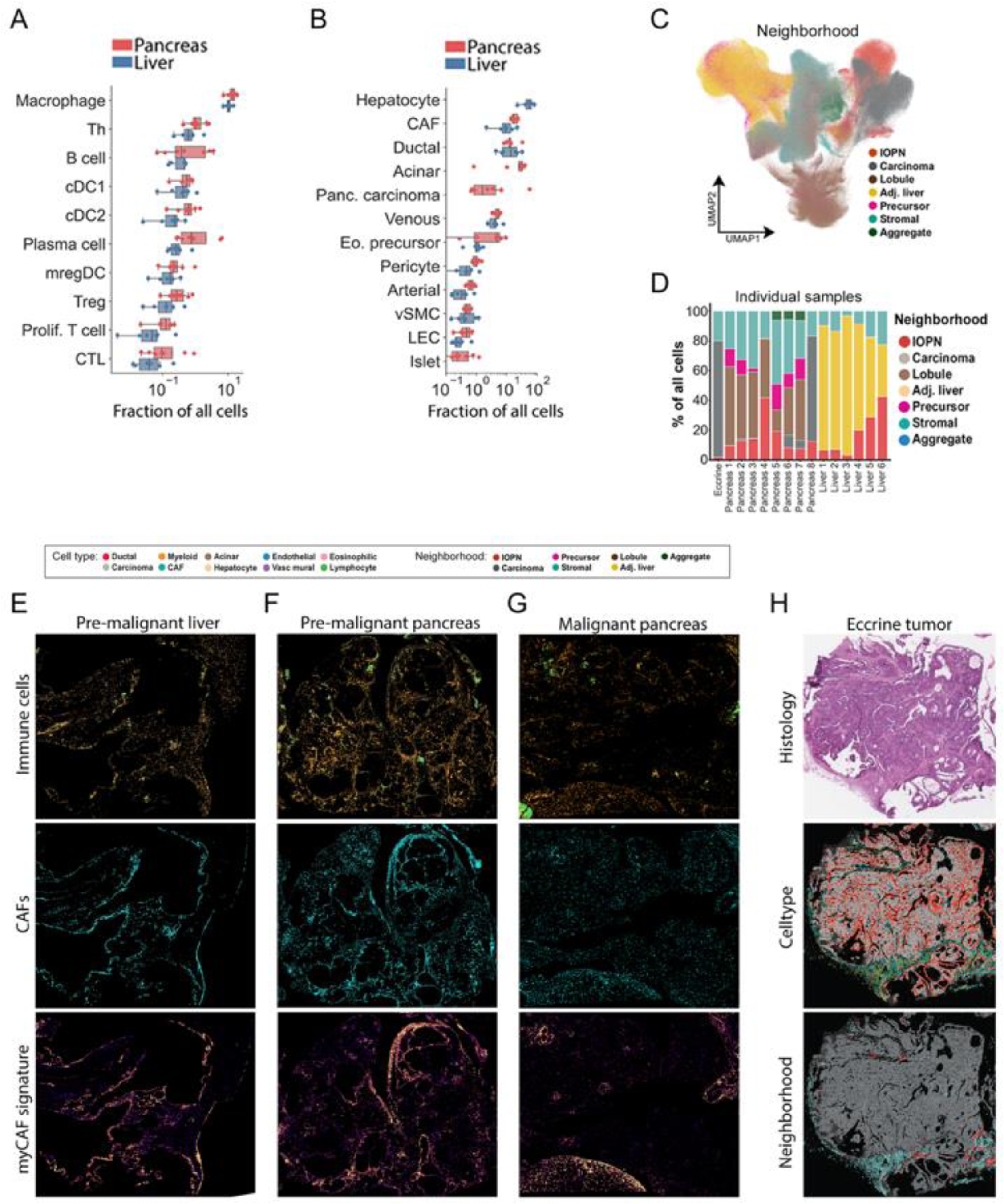
Spatial analysis of the tumor immune microenvironment in fusion driven tumors. **(A)** Relative immune cell frequencies are depicted for the pancreas and liver samples and demonstrate similar frequencies across these organs systems, with relative abundance of macrophages and paucity of cytotoxic T cells. **(B)** Relative cell type frequencies are depicted for pancreas and liver samples which demonstrate similar frequencies across organ systems other than expected tissue specific differences (i.e., hepatocytes in liver, islet cells in pancreas, etc.). **(C)** Uniform Manifold Approximation and Projection (UMAP) projection of 1,390,407 individual cells representing 15 mouse samples colored by cellular neighborhood. **(D)** Cell neighborhood frequencies from the spatial transcriptomics samples across individual samples. With the exception of tissue-specific populations (acinar, hepatocyte, islet), no significant differences were found across disease sites. **(E-G)** Immune cell, CAF, and myofibroblastic cancer associated fibroblast (myCAF) signature expression corresponding to section shown in Figure 6G-I. **(H)** Histology, cell type, and neighborhood for soft tissue tumor sample.

**Supplemental Table 1. Selected Gene Expression**

**Supplemental Table 2. Custom Gene Panel Xenium**

**Supplemental Table 3. Nucleotide Sequences**

